# B cells sustain inflammation and predict response to immune checkpoint blockade in human melanoma

**DOI:** 10.1101/478735

**Authors:** Johannes Griss, Wolfgang Bauer, Christine Wagner, Margarita Maurer-Granofszky, Martin Simon, Minyi Chen, Peter Steinberger, Katharina Grabmeier-Pfistershammer, Florian Roka, Thomas Penz, Christoph Bock, Gao Zhang, Meenhard Herlyn, Katharina Glatz, Heinz Läubli, Kirsten D Mertz, Peter Petzelbauer, Thomas Wiesner, Markus Hartl, Winfried F Pickl, Rajasekharan Somasundaram, Stephan N Wagner

**Affiliations:** Department of Dermatology, Medical University of Vienna, 1090 Vienna, Austria; EMBL-European Bioinformatics Institute, Wellcome Trust Genome Campus, CB10 1SD Hinxton, Cambridge, United Kingdom; Division of Immune Receptors and T-cell Activation, Institute of Immunology, Center for Pathophysiology, Infectiology and Immunology, Medical University of Vienna; CeMM Research Center for Molecular Medicine of the Austrian Academy of Sciences, 1090 Vienna, Austria; Department of Laboratory Medicine, Medical University of Vienna, 1090 Vienna, Austria; Molecular & Cellular Oncogenesis Program and Melanoma Research Center, The Wistar Institute, Philadelphia, PA 19104-4265, USA; Institute of Pathology, University Hospital Basel, Basel, Switzerland, 4031; Division of Medical Oncology, University Hospital Basel, Basel, Switzerland, 4031; Institute of Pathology, Cantonal Hospital Baselland, Liestal, Switzerland, 4410; Mass Spectrometry Facility, Max F. Perutz Laboratories (MFPL), University of Vienna, Vienna BioCenter (VBC), 1030 Vienna, Austria; Division of Cellular Immunology and Immunohematology, Institute of Immunology, Center for Pathophysiology, Infectiology and Immunology, Medical University of Vienna

## Abstract

Tumor associated inflammation predicts response to immune checkpoint blockade in human melanoma. Established mechanisms that underlie therapy response and resistance center on anti-tumor T cell responses.

Here we show that tumor-associated B cells are vital to tumor associated inflammation. Autologous B cells were directly induced by melanoma conditioned medium, expressed pro- and anti-inflammatory factors, and differentiated towards a plasmablast-like phenotype *in vitro*. We could identify this phenotype as a distinct cluster of B cells in an independent public single-cell RNA-seq dataset from melanoma tumors. There, plasmablast-like tumor-associated B cells showed expression of CD8+T cell-recruiting chemokines such as CCL3, CCL4, CCL5 and CCL28. Depletion of tumor associated B cells in metastatic melanoma patients by anti-CD20 immunotherapy decreased overall inflammation and CD8+T cell numbers in the human melanoma TME. Conversely, the frequency of plasmablast-like B cells in pretherapy melanoma samples predicted response and survival to immune checkpoint blockade in two independent cohorts. Tumor-associated B cells therefore orchestrate and sustain tumor inflammation, recruit CD8+ T effector cells and may represent a predictor for response and survival to immune checkpoint blockade in human melanoma.

---

Cancers such as melanoma, lung, and kidney cancer often present with an inflamed but immunosuppressive tumor microenvironment (TME). Immune checkpoint blocking (ICB) antibodies have significantly improved cancer therapy by overcoming inhibition of T cell effector functions. Yet, a considerable number of patients does not benefit from ICB therapy^1^. It is therefore key to understand the mechanisms that regulate inflammation within the TME to develop novel therapies and improve patient survival.

B cells promote both acute immune-associated inflammation for protection against foreign pathogens as well as chronic inflammation in autoimmune diseases and persistent infection. Mouse cancer models show that tumor-associated B cells (TAB) promote tumor inflammation^2,3^ but may also inhibit anti-tumor T cell-dependent therapy responses^4–7^. The immuno-inhibitory function of TAB in these models resembles that of regulatory B cells (Breg), which are an established source of inhibitory cytokines such as IL-10 and TGF-b (reviewed in ^8^). In human cancer, Breg were described by either phenotyping, direct detection of immunoinhibitory cytokines or surface molecules, and/or immunosuppressive function^4,9–13^. Often Breg frequencies increase with tumor progression and are enriched in tumors compared to peripheral blood or adjacent normal tissue. Increased IL-10^+^ B cell numbers can also be accompanied by increased numbers of CD4^+^CD25^+/high^CD127^low/-^ and Foxp3^+^ Tregs in tumor tissues^10,12,14,15^ which were independently associated with tumor progression or reduced patient survival.

In human melanoma, up to 33% of the immune cells can be TAB^16,17^ and phenotypic analysis has revealed CD20+ TAB (reviewed in^18^) and CD138^+^ or IgA^+^CD138^+^ plasma cells^17,19^. Conclusions about their impact on disease progression and outcome are inconsistent. So far, no data exist for TAB functions in murine models of melanoma highlighting the need for studies in melanoma patients and tumor samples. We recently showed that TAB-derived IGF-1 is a source of acquired drug resistance of human melanoma to mitogen-activated protein kinase (MAPK) inhibitors^16^. Clinical data from our pilot trial and an independent case series indicate objective tumor responses and clinical benefit through B cell-depletion by anti-CD20 antibodies in end-stage therapy-resistant metastatic melanoma patients^16,20^.

Human melanoma cells foster the generation of TAB with regulatory activity^21–23^. They provide antigens for prolonged B cell receptor stimulation and release inflammation-modulating cytokines such as IL-1b, IL-6^24^ and IL-35^25^. The functional phenotypes of TAB in human melanoma and their impact on inflammation and response to ICB therapy are, however, still unknown.

We therefore systematically characterized the effect of human melanoma cells on autologous B cells *in vitro* (Supplementary Figure 1). This induced a phenotypic and functional change of B cells towards a plasmablast-like phenotype with expression of pro- and anti-inflammatory molecules. This phenotype could be recapitulated in independent single-cell (sc) RNA-sequencing (seq) data from metastatic melanoma patients. In this data, plasmablast-like TAB also expressed several chemokines attracting immune cells, for example, T (reg) cells and macrophages. Consistently, depletion of TAB in metastatic melanoma patients drastically decreased overall inflammation and immune cell numbers in the TME and, conversely, the frequency of plasmablast-like TAB in pretherapy tumor samples was predictive for response and survival to ICB therapy.

## Results

### Melanoma TAB are composed of multiple phenotypes

We used 7 color multiplex immunostaining (CD20, CD19, CD5, CD27, CD38, CD138, and DAPI) to approximate 6 different TAB-subpopulations (Figure 1A, Supplementary Figure 2A) in tissue microarrays composed of 155 cores from human metastatic melanomas (Supplementary Figure 2B). In total, we observed 1,991 (45%) plasmablast-like TAB (Figure 1B), 1,089 (24%) plasma cell-like TAB (Supplementary Figure 3), 440 (10%) activated B cell-like TAB, 327 (7%) memory B cell-like TAB, 321 (7%) germinal-center B cell-like TAB, and 294 (7%) transitional B cell-like TAB. There was considerable variation between the cores (Supplementary Figure 2B). TAB are primarily located at the invasive tumor-stroma margin^16, 17, 26^. This suggests a preferentially contact independent communication between melanoma cells and TAB.

**Figure 1:**
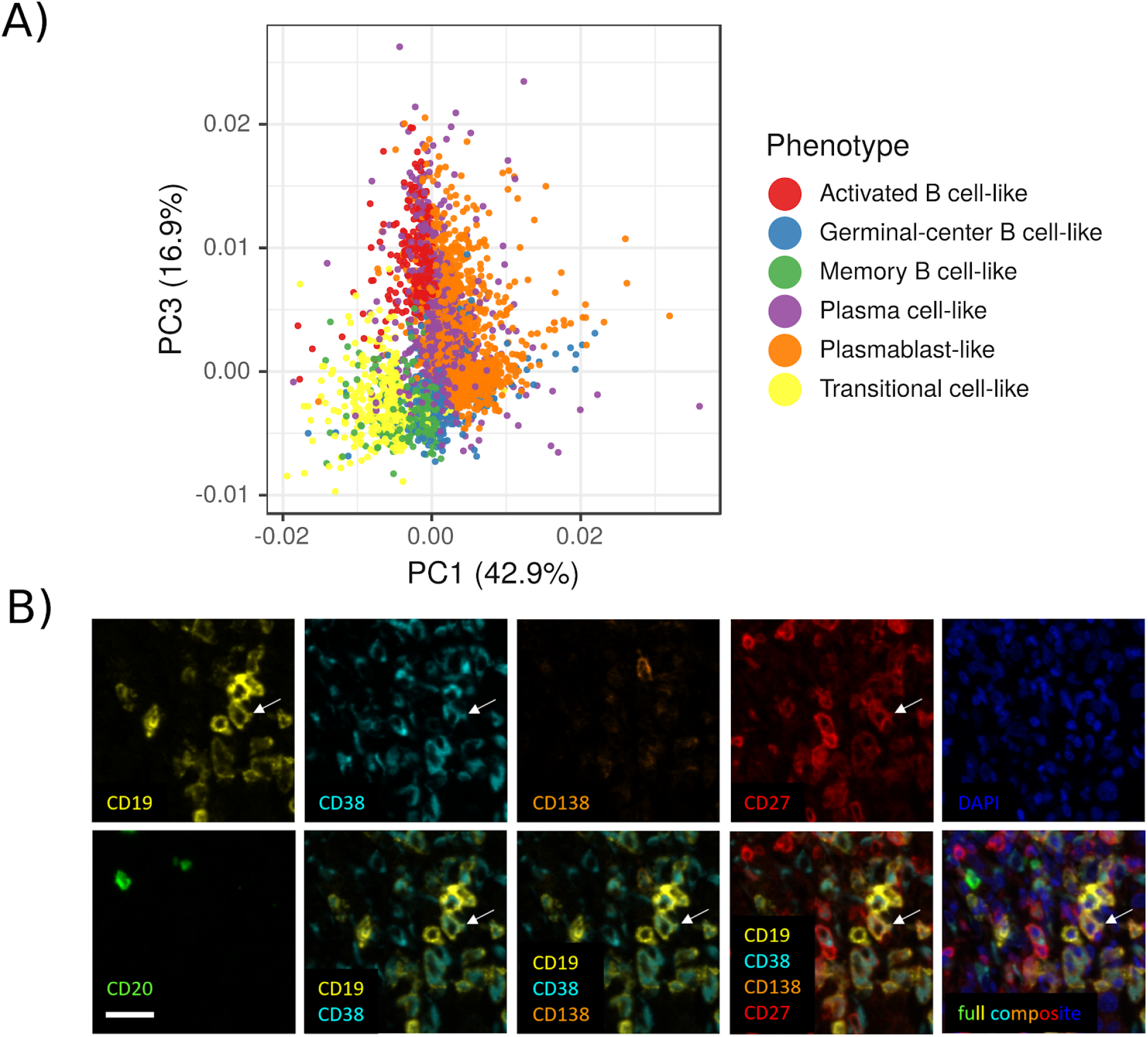
Melanoma TAB predominantly consist of plasmablast-like and plasma cell-like phenotypes. A) Principal component analysis of all characterized cells and their phenotypes in the 155 tissue microarray cores. The shown percentage represents the explained variance per component. Principal component analysis was performed on all staining intensities. B) Multiplex immunostaining of human melanoma identifies CD19+CD20-CD38+CD138-CD27+ plasmablast-like TAB. For each panel, serial images display the same cells from a stromal area at the invasive tumor margin. Composite image together with DAPI nuclear staining (bottom right) and images for each of the individual markers and different combinations from the composite image. Arrow depicts one of several plasmablast-like TAB. Scale bar represents 20μm.

### Human melanoma cells directly induce NFKB activation in TAB through soluble factors

The known release of pro-and anti-inflammatory cytokines from melanoma cells and our multiplex immunostaining results suggest that melanoma cells communicate through soluble factors with TAB. We therefore exposed immortalized peripheral blood- and tumor-derived B cells derived from 4 patients with metastatic melanoma to melanoma-conditioned medium collected from autologous early passage melanoma cells. In a screen for expression of the key immunoregulatory and pro-inflammatory cytokines/molecules PD-L1 and IL-6, all melanoma conditioned media induced a similar up-regulation in matched peripheral blood- and tumor-derived B cells (Supplementary Figure 4A). We therefore performed subsequent experiments with the melanoma conditioned medium from patient 2 as it induced the most prominent changes.

572 genes/proteins were identified as significantly regulated by melanoma conditioned medium in both, the RNA-seq and proteomics results. Additionally, 974 proteins were found only significantly regulated in the proteomics data and 2,520 genes found only significantly regulated in the RNA-seq data (Benjamini-Hochberg (BH) corrected p-value < 0.05, Supplementary Table 1). There was no marked difference between peripheral and tumor-derived B cells (Figure 2A). The estimated fold changes of genes identified as significantly regulated in both approaches showed a high linear correlation (Spearman cor = 0.79, p < 0.01, Supplementary Figure 4B). The subsequent pathway analysis (see Methods) showed upregulation of pathways associated with inflammation, immunity, B cell receptor signaling and intracellular signal transduction, and downregulation of pathways associated with cell cycle, cell division, DNA replication, DNA repair, translation, and transcription (Supplementary Table 1).

**Figure 2:**
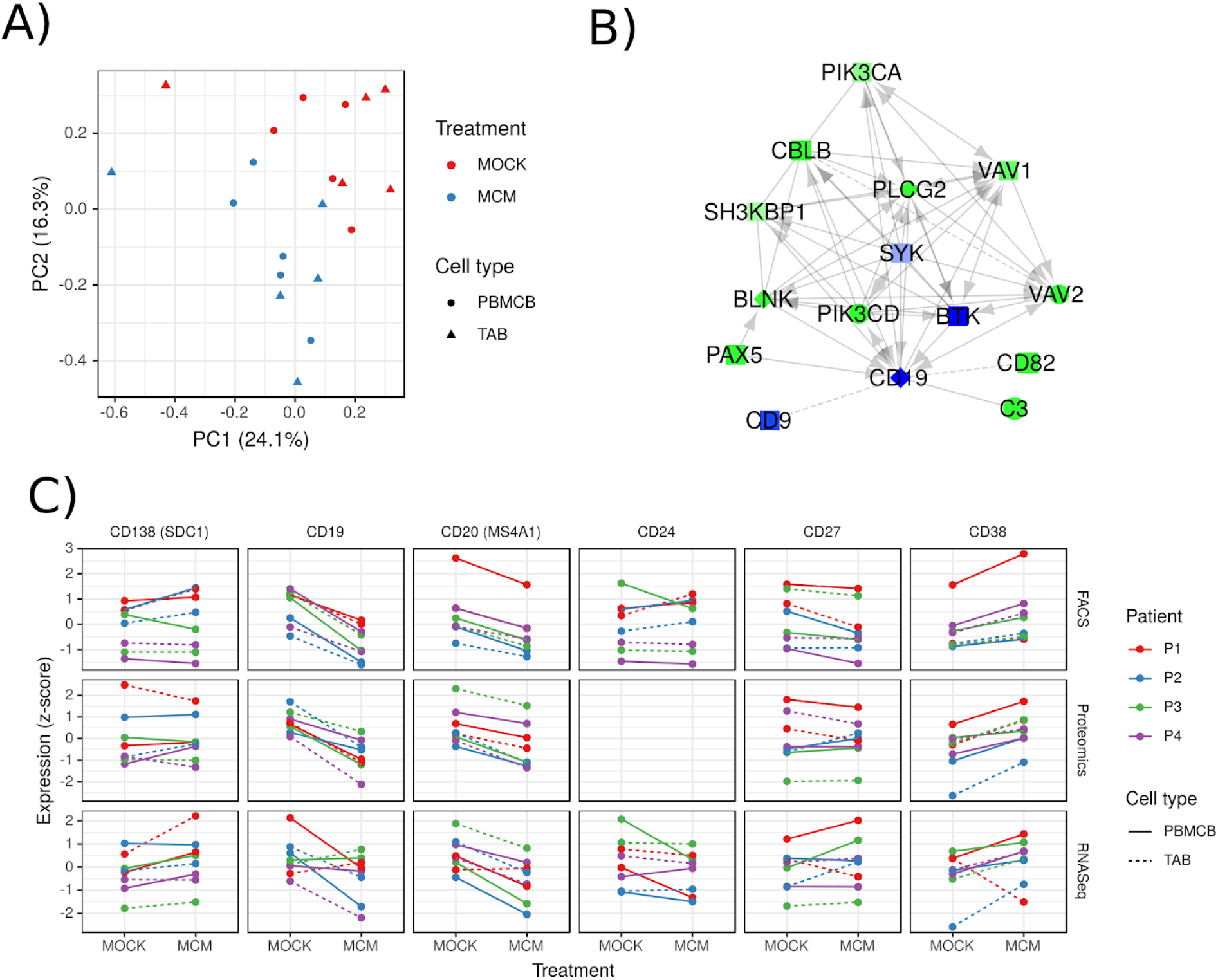
Melanoma conditioned medium induced distinct phenotypic changes in TAB. A) Principal component analysis of all samples based on the proteomics data comparing samples treated with melanoma conditioned medium (MCM) against control treated ones (MOCK). Peripheral blood derived B cells (PBMCB) and TAB are shown through different shapes. The percentage in the axis labels represents the variation explained by the component. Points represent one sample per patient and one sample where individual patient samples were pooled together. B) Interaction network of CD19 with the nearest neighbours depicting the gene abundance estimated by RNA-seq and proteomics. Green represents up-regulation, blue down-regulation, light colors indicate no relevant difference. Diamonds are genes only detected in proteomics, rectangles genes only detected in RNA-seq, circles genes identified by both methods. For the actual data analysis, the network was constructed using the two nearest neighbours. C) Expression of key differentiation markers estimated by FACS (geo mean expression), proteomics and RNA-seq for melanoma conditioned medium (MCM) and control (MOCK) treated immortalized peripheral blood derived B cells (PBMCB) and TAB. All values are shown as z-scores.

One of the most significantly upregulated pathways was “tumor necrosis factor (TNF) signalling via NFKB”. NFKB is a key transcription factor for B cell activation in inflammation and immune response^27^. This pathway includes, among others, CD69, CD80, CD30 (TNFRSF8), and CD137 (4-1BB/TNFRSF9) which were significantly upregulated in our proteomics and transcriptomics data (with CD30 (TNFRSF8) among the top-20 upregulated genes) and are known to be upregulated in B cells upon activation. Even though TNF itself showed no significant differences, TNF alpha induced protein 2 (TNFAIP2) was significantly upregulated as indirect evidence for TNF signalling. In addition, we observed the up-regulation of CD40 signalling genes which also acts via NFKB. We also found increased phosphorylation of PRKCB which plays a key role in B cell activation by regulating BCR-induced NFKB activation (with no significant differences in transcriptomics or global proteomics analysis). Similarly, we observed an increased phosphorylation of NKAP which is involved in TNF- and IL-1 induced NFKB activation. Therefore, melanoma-derived soluble factors directly induce a signalling pattern in B cells associated with activation in inflammation and immune response.

### Melanoma conditioned medium induces a plasmablast-like dominated TAB population

The downregulation of cell cycle associated pathways coincided with a significant, unexpected down-regulation of CD20 (MS4A1, BH adjusted p < 0.01 RNA-seq) and CD19 (BH adjusted p < 0.01 proteomics). This indicates a phenotypic change of B cells. To identify this phenotype, we first used the Reactome functional interaction network^28^ and extracted all neighbours of CD19 and their neighbours (Figure 2B). While known key developmental phenotypes, including reported regulatory B cell markers such as CD147, CD24, CD27, CD25, CD39, CD73, or CD138 together with transcription factors BLIMP-1, XBP-1 were not significantly regulated, downregulated (IRF4, CD71) or not detectable (CD5, TIM1), we found a significant upregulation of CD38 (proteomics, BH adjusted p = 0.01), together with PAX5 (BH adjusted p < 0.01 RNA-seq).

Secondly, 9-color FACS staining showed changes of CD19, CD20, CD24, CD27, CD38 and CD138 consistent with proteomics and RNA-seq results (Figure 2C). Immunoglobulin D, M and G expression was unchanged (Supplementary Figure 4C). The induction of CD38 points towards a plasmablast/–cytoid differentiation. This is counter-regulated by increased PAX5 expression. These findings correlate with the high proportion of plasmablast-like TAB in human melanomas (Supplementary Figure 2B). Based on these results, we defined a tumor-induced plasmablast-like dominated B cell population (TIPB) signature with the genes CD27, CD38, and PAX5.

### Melanoma conditioned medium induces distinct functional signatures in TAB which are found *in vivo*

To profile TIPB on a functional level, we manually extracted six key immunological functional gene sets from the melanoma conditioned medium induced, enriched CD19 interaction network (Figure 2B, see Methods). To reduce the bias invariably introduced by our small sample size, we additionally extracted highly correlating genes from the TCGA skin cutaneous melanoma dataset to create six signatures based on existing literature. In line with the pathway analysis (increased signalling via NFKB), these key upregulated gene sets included *activation-associated genes* (CD69, CD72, CFLAR, FGFR1, SELPLG, CD86), *co-stimulatory genes* (ICAM3, TNFRSF13C, CD40, CD72, C3, CD80, CD86, CD27, CD28, ICOS, TNFRSF9, CD40LG, ARHGDIB), and *pro-inflammatory genes* (TNF, IL12B, IL18, LTA, TNFAIP2, C3, HCK). In addition, *immune checkpoint-associated genes* (CD274, PDCD1LG2, TNFRSF14/HVEM, LGALS9, BTLA, LAG3, HAVCR2/TIM-3, ADORA2A), *immunosuppressive genes* (IL10, TGFB1), as well as *B cell exhaustion-associated genes* (PDCD1, FCRL4, SIGLEC6, CD22) were upregulated.

We used scRNA-seq data by Sade-Feldman *et al*. on immune-checkpoint blockade and response in human melanoma^29^ to validate these functional signatures. TAB clustered into distinct groups which we correlated with known developmental B cell phenotypes (Figure 3A). Overall, all B cell phenotypes expressed comparable levels of our functional signatures (Figure 3B) but with distinct differences on the single gene level (Supplementary Table 2). Plasmablast-like and memory-like B cells expressed substantial levels of immunosuppressive cytokines such as TGFB1 and IL-10 (Figure 3C). We could additionally validate the expression of the latter through multiplex immunostaining in plasmablast-like B cells (Figure 3D). Remarkably, TAB also expressed numerous chemokines including T (reg) cell, and macrophage chemoattractants CCL3, CCL3L1, CCL4, CCL5, CCL28 and CXCL16^30–33^ in plasmablast-like B cells (Figure 3C). Together, these results indicate that TIPB are able to regulate inflammation and shape the cellular composition of the human melanoma TME.

**Figure 3:**
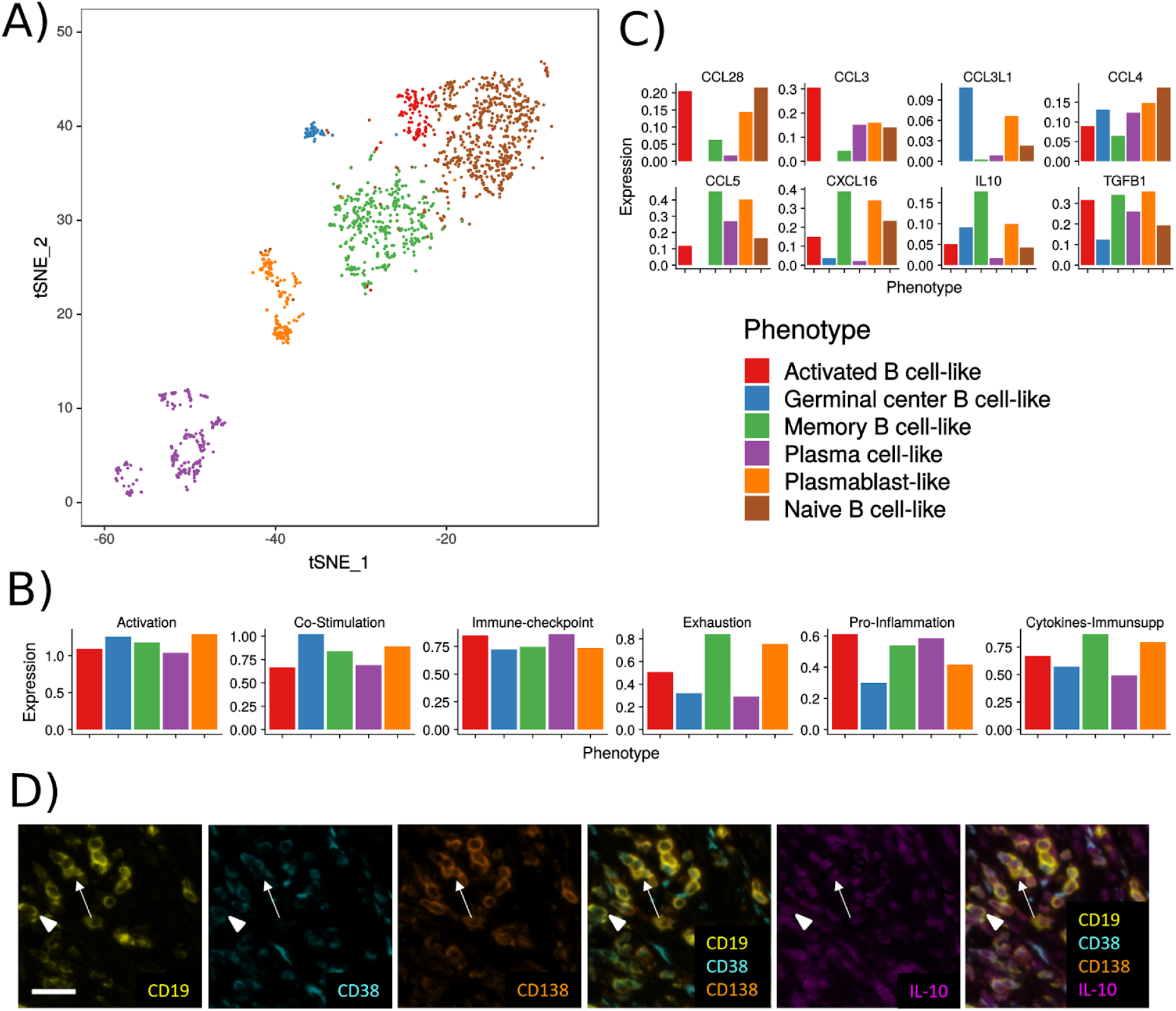
scRNA-seq data from human melanoma verifies the observed functional phenotypes *in vivo*. A) T distributed stochastic neighbor embedding representation of all B cells with their respective phenotype annotations from the Sade-Feldman *et al*. dataset. B) ssGSEA estimated expression of the functional signatures based on the average gene expression per B cell phenotype. C) Average expression of specific chemokines and immunosuppressive cytokines per B cell phenotype. D) Confirmation of IL-10 expression in plasma cell-like (arrow) and plasmablast-like (arrowhead) TAB by quadruple marker immunostaining. Scale bar represents 20 μm.

### TIPB correlate with the predicted functional signatures, CD8+ T cells, macrophages, and overall survival

Next, we tested whether the TIPB signature correlates with the identified functional signatures in independent melanoma cohorts and evaluated its effect on clinical outcome in the TCGA skin cutaneous melanoma cohort. All functional signatures (except for the *immunosuppressive genes*, Spearman correlation 0.49, BH adjusted p < 0.01) showed a strong linear correlation with our TIPB signature (Spearman correlation >= 0.78, BH adjusted p < 0.01, Supplementary Figure 5A). Our TIPB signature was also significantly correlated with the expression of CD8A (Spearman correlation 0.87, p < 0.01, Supplementary Figure 5B) as well as with the abundance of CD8+ T cells (Spearman correlation 0.82, p < 0.01, Supplementary Figure 5C) and macrophages (Spearman correlation 0.71, p < 0.01) as estimated by xCell^34^. Established signatures describing the inflammatory TME and possible response to anti-PD1 therapy (tumor inflammatory score^35^, interferon gamma^36^, T-cell exhaustion^37^, and T-cell effector^38^ signatures) additionally showed a high linear correlation with our TIPB signature (Spearman correlation >= 0.85, BH adjusted p < 0.01, Supplementary Figure 5D). Finally, high expression (above median) of our TIPB signature was correlated with longer overall survival in the TCGA cohort (Figure 4A). These results show, that the TIPB and the predicted functional signatures, are highly associated with T cell abundance and inflammation, and, importantly, are linked with patient outcome.

**Figure 4:**
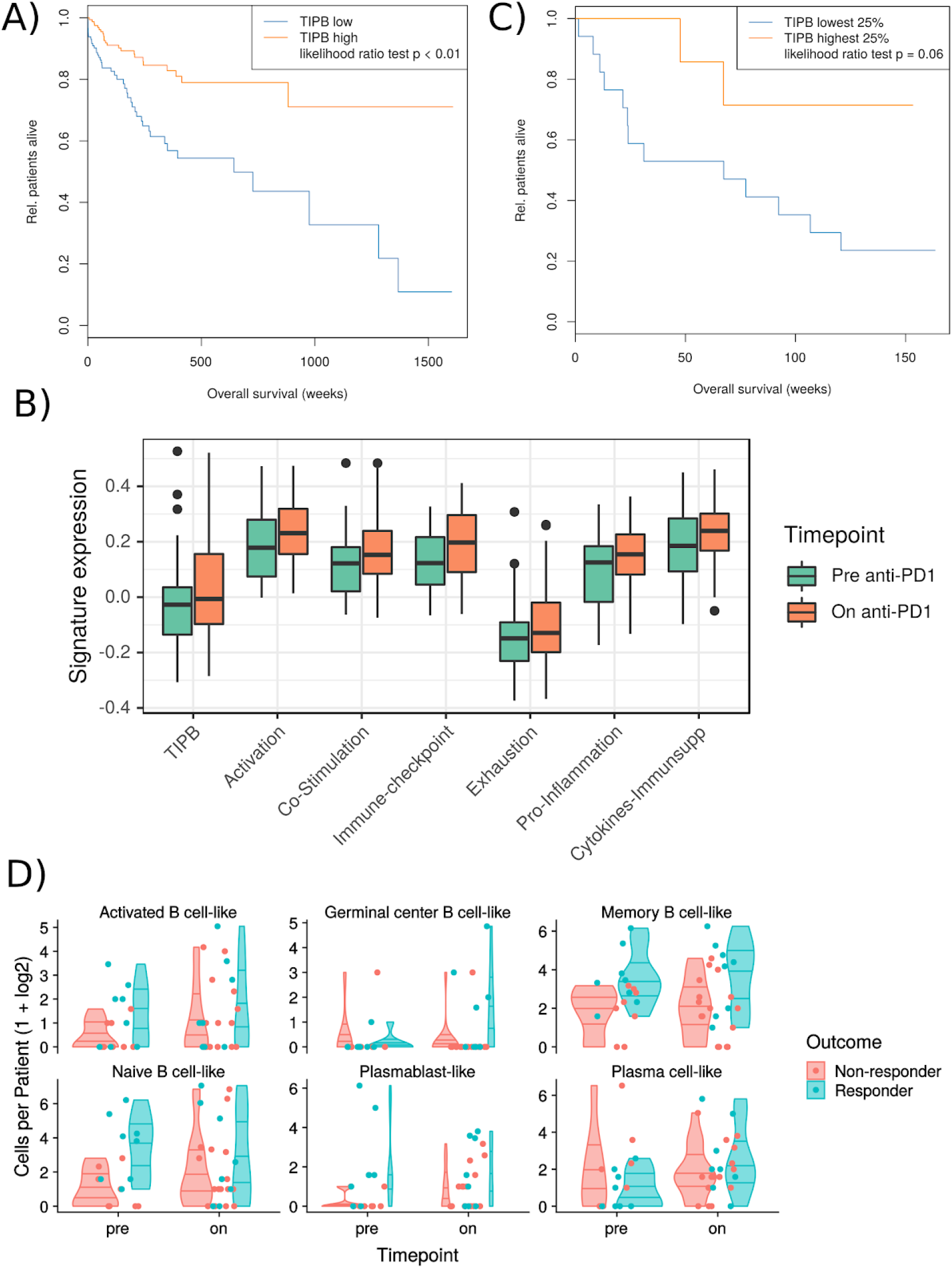
TIPB predict improved survival of melanoma patients. A) Survival analysis of patients expressing high or low levels of the TIPB signature separated by the median expression in the TCGA melanoma cohort. B) Expression of the TIPB signature and all functional signatures before and on anti-PD1 therapy as estimated in the Riaz *et al*. dataset. C) Survival analysis based on the TIPB signature in the pre-treatment samples comparing top 25% expressing samples against the lower 25% samples in the Riaz *et al*. dataset. D) Frequencies of B cell phenotypes (logarithmic scale) before and on ICB therapy in the scRNA-seq dataset from Sade-Feldman *et al*. separated by response. Plots represent the relative frequencies, lines represent the 25%, 50%, and 75% quantiles.

Anti-PD1 therapy frequently leads to an increase in B cell numbers which should enhance our functional signatures. We used the transcriptomics data by Riaz *et al*. containing (partially matched) 51 pre-anti-PD1 therapy and 58 post-anti-PD1 therapy samples^39^. In this independent cohort, all signatures with exception of the immunosuppressive genes (Spearman correlation = 0.6, BH adjusted p < 0.01) showed a strong linear correlation with the TIPB signature (Spearman correlation >= 0.77, BH adjusted p < 0.01, Supplementary Figure 6A). Again, the TIPB signature correlated with CD8A expression (Spearman correlation 0.9, BH adjusted p < 0.01, Supplementary Figure 6B), estimated CD8+ T cell (Spearman correlation 0.76, p < 0.01, Supplementary Figure 6C) and macrophages (Spearman correlation 0.68, p < 0.01) abundance. We also observed a consistent up-regulation of all signatures during anti-PD1 therapy (one-sided t-test BH adjusted p <= 0.03, df = 100-107, t = −2.03 - −2.77, immunosuppressive genes BH adjusted p = 0.06, df = 96.33, t = −1.58, exhaustion BH adjusted p = 0.09, df = 106.7, t = −1.35, Figure 4B). Interestingly, high expression of our TIPB signature before therapy (top 25% versus lower 25%) predicted overall survival (Likelihood ratio test p = 0.06, Figure 4C). While the TIPB signature did not correlate with clinical response (Supplementary Figure 7) in this dataset, scRNA-seq showed plasmablast-like and naive-like B cell frequencies in pre-therapy tumor samples to be significantly higher in patients responding to ICB therapy (two-sided Wilcoxon rank sum test, BH adjusted p-value = 0.04, Figure 4D). Together, these data validate the clinical importance of the identified TIPB population.

### Loss of TAB reduces inflammation in the tumor microenvironment

Finally, we evaluated the loss of TAB in a cohort of patients with metastatic melanoma treated with anti-CD20 antibodies^16, 39^ (see Methods, Supplementary Figure 1). The dataset consists of 4 patients with pre- and on-therapy samples of established metastases (affected by anti-CD20 therapy, therapeutic setting) and 2 patients with pre- and on-therapy samples, where the metastases developed *de-novo* in B cell depleted patients on therapy^40^ (adjuvant setting). Principal component analysis of whole tissue RNA-seq showed no systematic difference between the two patient groups (Figure 5A).

**Figure 5:**
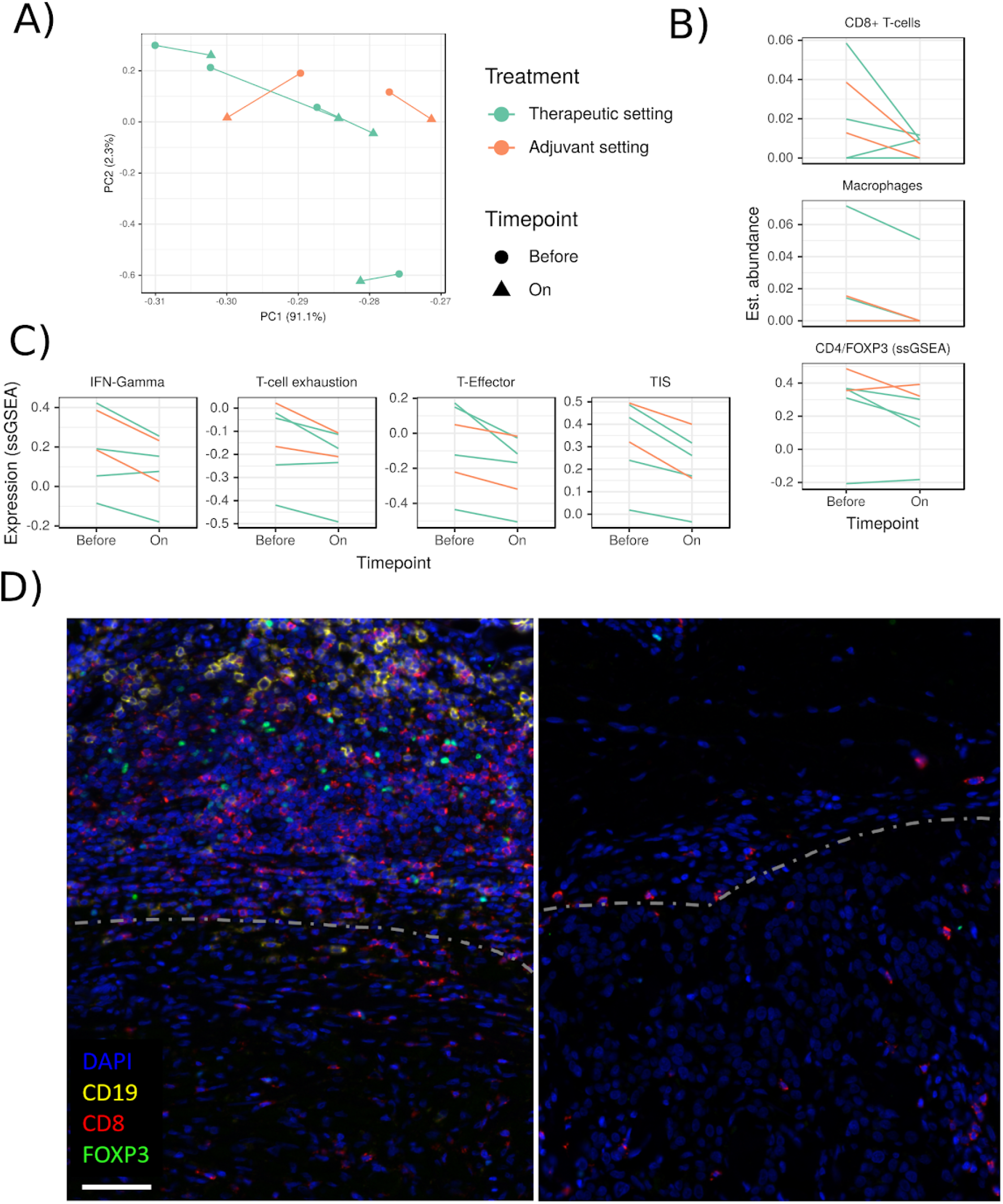
Reduction of TIPB through anti-CD20 therapy reduces tumor inflammation and CD8+ T cell numbers. A) Principal component analysis of RNA-seq data from samples before and on anti-CD20 therapy. On-therapy samples consist of metastases affected by anti-CD20 therapy (therapeutic setting, green lines) and of metastases that developed *de-novo* in B cell depleted patients (adjuvant setting, orange lines). Percentage numbers in axis labels represent the explained variation by each component. Lines link a patient’s samples. B) xCell estimated abundance of cell types in tissue samples before and on anti-CD20 therapy. Abundance of CD4/FOXP3 was estimated using ssGSEA since no comparable xCell signature exists. C) Expression of established inflammation (interferon (IFN) gamma, tumor inflammatory score (TIS) and T cell gene signatures before and on anti-CD20 therapy. D) Triple marker immunostaining of a representative patient-matched pair of melanoma samples before (left) and on (right) anti-CD20 therapy. Images were taken at the invasive tumor (bottom)-stromal tissue (upper part) margin (dashed line). Note disappearance of CD19 cells, and strong reduction of CD8+ and FoxP3+ cells after therapy. Scale bar represents 60 μm.

Next to the expected down-regulation of CD19 and CD20 (MS4A1), all patients showed a consistent, significant down-regulation of CD8A on anti-CD20 therapy (BH adjusted p = 0.02, Supplementary Table 3). This was linked to a consistent reduction of CD8+ T-cells (one-sided, paired t-test, BH adjusted p = 0.07, t = 1.74, df = 5) and macrophages (one-sided, paired t-test, BH adjusted p = 0.07, t = 2.17, df = 5, Figure 5B). A signature consisting of CD4 and FOXP3 to approximate the abundance of FOXP3+ T cells also decreased significantly (one-sided, paired t-test p = 0.05, df = 5, t = 2.03, Figure 5B).

Anti-CD20 therapy caused a significant downregulation of our TIPB signature and all our functional signatures (one sided, paired t-test, BH adjusted p-value 0.01 - 0.04, t = 2.3 - 4.8, df = 5, Supplementary Figure 8A). Published signatures describing the inflammatory TME and possible response to anti-PD1 therapy (tumor inflammatory score^35^, interferon gamma^36^, T-cell exhaustion^37^, and T-cell effector^38^ signatures) highly correlated with our TIPB signature (Spearman correlation 0.7 - 0.9, BH adjusted p < 0.01, Supplementary Figure 8B) and showed a significant decrease on anti-CD20 therapy (one-sided, paired t-test, BH adjusted p = 0.01, df = 5, t = 3.11 - 5.46, Figure 5D). These data validate the essential role of TIPB to sustain tumor inflammation and recruit CD8+ T cells and macrophages in human melanoma.

## Discussion

Tumor-associated inflammation classifies human melanoma into two major immunological subtypes, T cell-inflamed or non-inflamed (immune-excluded or -desert), with different response to therapy and patient prognosis^41^. Our data show that TAB profoundly modify and sustain inflammation through melanoma-induced subpopulations enriched for plasmablast-like cells, that we called TIPB, with co-expression of pro- and anti-inflammatory factors. These data could be confirmed in most recent independent public scRNA-seq data, which showed that different B cell subtypes including plasmablast-like cells co-express our identified functional signatures.

The profound effects of TIPB and their associated functional signatures in human melanoma are best demonstrated by TAB depletion therapy, which caused a shift towards a less inflamed TME with distinct loss of CD8+ T cells and macrophages. These changes have important clinical implications as they are linked with resistance to MAPKi therapy^42^ and with established signatures predictive of survival and response to PD-1 immunotherapy (reviewed in ^43^). Conversely and most importantly, the frequency of TIPB in pre-therapy tumor samples was associated with response^29^ and overall survival^39^ of patients undergoing PD-1 immune checkpoint blockade therapy. In view of the present lack of predictive markers, this association bears enormous potential clinical and socioeconomic implications.

The co-expression of immunostimulatory and -inhibitory cell surface receptors characterizes TIPB as a rather unique B cell population. Melanoma-induced stimulatory signatures indicated antibody-independent functions of TIPB, such as antigen presentation, T cell activation and cytokine production with the ability to promote local inflammation as recently shown in multiple sclerosis^44^. TIPB also expressed inhibitory receptors and ligands as well as IL10 and TGFB1 as indicated by scRNA-seq and multiplex immunostaining data. Therefore, TIPB resemble Breg, which directly and indirectly suppress proliferation, anti-tumor activity, and differentiation of several immune cells residing in the TME^45–50^. The expression of these inhibitory factors in TIPB is in line with previous reports on human CD27^int^CD38^+^CD138^-^ plasmablasts as a major IL-10+ B cell population^47^, human colorectal cancers being highly enriched for IL-10+CD19^lo^CD27^hi^ plasmablasts that can suppress IL-17A^+^CD4^+^ T cells^51^ and murine IgA^+^PD-L1^+^L10^+^ plasmocytes that prevent CD8^+^ T cell-mediated regression of hepatocellular carcinoma^7^ and impede T cell-dependent chemotherapy of castration-resistant prostate cancer^4^. Our data expand these observations by pinpointing the additional expression of inhibitory ligands such as LGALS9/Galectin-9, and TNFRSF14/HVEM in TIPB. These ligands can stimulate their cognate inhibitory receptors HAVCR2/TIM-3 and BTLA, which are induced on T cells upon persistent antigen exposure and drive T cell dysfunction.

The “tide model” describes expression of co-stimulatory and inhibitory signals as a dynamic system to fine-tune immune responses^52^. Additionally, the “immune set point model” highlights the often subtle contribution of intrinsic and extrinsic factors that modulate tumor inflammation and tip the balance between immunity and tolerance^41^. In human melanoma, even small variations within the release of tumor antigens/neoantigens, extrinsic or environmental factors, the secretion and consumption of cytokines and chemokines as well as therapeutic agents can tip this balance. The well-known dichotomy of clinical responses to ICB could thus reflect the overall inflammatory status of the TME^41^. Similarly, pro-tumorigenic^16^ as well as pro-immunogenic functions of TIPB can result from a disbalanced expression of inflammation-modulating factors and depend on the degree of inflammation in the respective TME.

The functional plasticity of TIPB emphasizes the importance of a careful evaluation of their functions in different tumor and host contexts to guide the development of TIPB-targeting strategies for novel cancer combination therapies.

## Acknowledgements

This work was supported by the FWF-Austrian Science Fund (project P31127-B28 to SNW and SFB F4609 to WFP) and received funding from the European Research Council (ERC) under the European Union’s Horizon 2020 research and innovation programme under grant agreement No 788042. We thank Bjoern Wendik (Perkin Elmer) for support in tissue imaging and data acquisition of multiplex immunostainings, Barbara Sternitzky and Claudia Kokesch for technical support in processing tissue microarrays, the Biomedical Sequencing Facility at CeMM for assistance with next-generation sequencing, the MFPL Mass Spectrometry team for technical support with proteomics, and the VBCF for providing the mass spectrometry instrument pool. This work was supported in part by NIH grants P01 CA114046, P50 CA174523, U54 CA224070, DoD - PRCRP WX1XWH-16-1-0119 [CA150619], and the Dr. Miriam and Sheldon G. Adelson Medical Research Foundation to MHe.

## Author contributions

**Hypothesis & study protocol** JG, GZ, MHe, RS, SNW **Clinical study & patient materials** JG, MMG, FR, PP, WFP, SNW **Histopathology** CW, MC, KG, HL, KDM, PP, SNW **FACS** WB, MS, PS, KGM **Proteomics** JG, MMG, MHa, CW **RNA-seq** JG, TW, CW, TP, CB **Bioinformatics** JG **Manuscript** JG, SNW.

All authors contributed to the final version of the manuscript.

## Methods

### Data availability

RNA-seq data were deposited in the ArrayExpress^53^ database at EMBL-EBI (www.ebi.ac.uk/arrayexpress) under accession numbers E-MTAB-7472 (induction experiment) and E-MTAB-7473 (anti-CD20 study). The mass spectrometry proteomics data were deposited to the ProteomeXchange Consortium via the PRIDE partner repository^54^ with the dataset identifier PXD011799 (induction experiment).

### Patient-derived material

Of 10 patients with metastatic melanoma who were treated with the anti-CD20 antibody ofatumumab in a therapeutic setting^16,55^(Clinical trials number NCT01376713), six met the selection criteria for inclusion into this analysis. Patients had to have pre-therapy peripheral blood mononuclear cell (PBMC) samples for generation of EBV-immortalized peripheral B cells as well as adequate pre-therapy biopsy material for generation of (*i*) early passage melanoma cells and (*ii*) autologous EBV-immortalized TAB from tumor single cell suspensions. From three patients these samples were successfully established. Or patients had to have adequate biopsy material for RNA-seq analysis at two time points: before initiation of anti-CD20 therapy and under therapy from week 8 on with biopsies taken under therapy showing a depletion of CD20+ TAB. From matched samples of four patients, RNA-seq data were successfully generated, including one patient from the former cohort. To increase the number of patients with pre-therapy early passage melanoma cells and autologous EBV-immortalized peripheral B cells and TAB, we included one further patient who was considered for but did not undergo anti-CD20 therapy.

In addition, of two patients with metastatic melanoma who were treated with the anti-CD20 antibody rituximab in an adjuvant setting^40^ (EudraCT number: 2007-005125-30), adequate tumor biopsy material for RNA-seq was available at two time points: before initiation of anti-CD20 therapy, when the tumor mass was completely resected, and after disease progression under therapy. Both studies were investigator-initiated clinical pilot trials. Anti-CD20 antibodies (Ofatumumab vs. Rituximab) were chosen based on availability. Samples were collected under local ethics committee-approved protocols (457-2007, 969-2010) with informed patient consent. Separate ethics committee-approval was obtained to make the derived proteomics and RNA-seq data publicly available (1555-2016). Clinical details are summarized in Supplementary Table 4.

### Induction experiments with Melanoma-conditioned medium (MCM)

Immediately after surgical resection, melanoma cells and TAB were collected by mechanical and enzymatic dissociation of metastatic melanoma tissue into single cells (tumor dissociation kit, gentleMACS Dissociator, both Miltenyi Biotec GmbH, Bergisch Gladbach, Germany) as described earlier by us^55,56^. Peripheral blood samples were collected in BD Vacutainer CPT cell preparation tubes and PBMC prepared according to the manufacturer’s instructions.

Melanoma cells grown from these single cell suspensions were frozen after passage 2 and stored in liquid nitrogen. Cell purity was determined by flow cytometric staining for melanoma-associated chondroitin sulfate proteoglycan (MCSP) on a FACS Calibur (Becton Dickinson, San Jose, CA). The percentage of MCSP-positive cells was >97%^56^. Sorting of TAB from single cell suspensions and peripheral B cells from PBMC was performed with Dynabeads CD19 Pan B (Life Technologies, Grand Island, NY) as described^55^. After detachment of beads and antibodies (DETACHaBEAD CD19 Kit, Thermo Fisher Scientific), flow cytometric analyses with a (PE)-conjugated CD19 antibody (Miltenyi Biotec) revealed a purity of collected B cell samples between 89.3 and 98.4%, respectively. Immortalization of TAB and peripheral B cells with EBV was performed on autologous irradiated (2× 30 Gy) PBMC as feeder cells essentially as described^55,57^. The purity of immortalized B cells was assessed as >96% in all samples by flow cytometry. A semi-quantitative comparative proteome of autologous freshly isolated vs. immortalized TAB has revealed significant differences predominantly in pathways related to cell cycle proliferation, apoptosis, and interferon response^55^. In addition, we have recently shown that immortalized TAB and peripheral blood-derived B cells can nicely recapitulate the induction of pro-tumorigenic/pro-inflammatory factors/cytokines in freshly isolated TAB and peripheral blood-derived B cells upon exposure to soluble factors from human melanoma cells^16,55^ and, thus, can offer a consistent source of quality cells for such experiments.

To produce MCM, early passage (p3) human melanoma cells were grown in complete RPMI 1640 medium (Life Technologies) supplemented with 10% FCS, 100 IU penicillin G, and 100 mg/ml streptomycin at 37°C in a humidified atmosphere with 5% CO_2_. At 80% confluence, the entire media was replaced with fresh medium. MCM was collected 48 hrs thereafter and sterile-filtrated. Control medium was prepared the same way, but without melanoma cells. For induction experiments, 5×10^5^ human B cells were seeded in 24 well cell culture plates (Corning Costar) and incubated in 1 ml MCM or control medium for 48 hrs. About 50% of media was substituted with fresh MCM after 24 hrs. B cells were then pelleted by centrifugation at 400*g* for 5 min at RT, snap-frozen and stored at −80 °C.

### FACS analysis

Mock- or MCM-treated immortalized B cells or TAB were stained with the following antibodies or matched isotypes and analyzed on a FACS Aria III (BD): CD19 BV711 (clone SJ25C1), CD20 AF700 (clone 2H7), CD24 PE-CF594 (clone ML5), CD27 BV421 (clone M-T271), CD38 APC (clone HIT2), CD138 PE (clone MI15), IgD PE-Cy7 (clone IA6-2), IgG FITC (clone G18-145), IgM BV605 (clone G20-127) (all BD biosciences). Live/dead cell exclusion was performed by addition of 7-AAD (5μg/ml, Calbiochem) prior to acquisition of the samples. Data were analysed using FlowJo 10.4.2 (FlowJo LLC). The gating strategy is shown in the Supplementary Methods.

### Proteomics analysis

#### Cell lysis and protein digestion

All reagents were of analytical grade and obtained from SIGMA-Aldrich, unless specified otherwise. Cells were lysed in freshly prepared lysis buffer containing 100 mM Tris/HCL pH 7.6, 2 % sodium dodecyl sulfate (SDS), 1 mM sodium vanadate, 1 mM NaF, protease inhibitor (cOmplete^Tm^ EDTA-free) and phosphatase inhibitor (PhosSTOP^Tm^) cocktail tablets (both Roche). Cell extraction and DNA sharing was assisted by sonication and cell debris pelleted by centrifugation at 20.000 x g for 15 min at 20°C. The supernatant was collected and the total protein concentration determined using the BCA protein assay kit (Pierce Biotechnology). Filter-aided sample preparation (FASP) was performed using Amicon Ultra Centrifugal 30 kDa molecular weight cutoff filters (Millipore) essentially according to the procedure described by^58^. Dithiothreitol (DTT) was added to a final concentration of 0.1 M and the samples heated at 99°C for 5 minutes. 200 μL of each protein extract was mixed with 3.8 ml_ of 8 M urea in 100 mM Tris-HCl, pH 8.5 (UA) in the filter unit and centrifuged at 4000 g for 30 min at 20°C to remove SDS. Any remaining SDS was exchanged by urea in a second washing step with 4 mL of UA. Free thiols were alkylated with 2 mL of 50 mM iodoacetamide for 30 min at RT. Afterwards, three washes with 3 mL of UA solution and then three washes with 3 mL of 50 mM triethylammonium bicarbonate (TEAB) were performed. Proteins were digested on filters with trypsin (1:50; Trypsin Gold, Promega) in 50 mM TEAB overnight at 37°C. Digested peptides were collected by centrifugation, acidified with trifluoroacetic acid (TFA), and desalted using Sep-Pak C18 SPE cartridges (50 mg, Waters Corporation) using 80 % acetonitrile containing 0.1 % TFA for the elution and evaporating the solvent in a vacuum centrifuge.

#### TMT 10plex-labeling of peptides

Isobaric labeling was performed using 10plex tandem mass tag (TMT) reagents (Thermo Fisher Scientific). 200 μg peptide digest per cell line was resuspended in 100 mM TEAB buffer and labeled with 0.8 mg TMT 10-plex™ reagents (Thermo Fisher Scientific) according to the manufacturer’s protocol. After one hour incubation at room temperature, samples were quenched for 15 min with 8 μl 5 % hydroxylamine at room temperature. Labeling efficiency was determined running aliquots of the samples on 1h LC-MS/MS gradients and standard database searches with TMT-tags configured as variable modifications. Corresponding TMT-labeled samples were pooled, acidified with TFA to a final concentration of 1 % TFA and concentrated via Sep-Pak C18 SPE cartridges (200 mg bed volume).

#### Off-line basic pH reversed-phase fractionation

Off-line high-pH reversed phase fractionations was performed essentially according to^59^ with a few modifications. Peptides were separated on a Waters Xbridge BEH130 C18 3.5 μm 4.6 x 250 mm column on an Ultimate 3000 HPLC (Dionex, Thermo Fisher Scientific) operating at 0.8 ml/ min. Buffer A consisted of H_2_O and buffer B consisted of 90 % acetonitrile (MeCN), both adjusted to pH 10 with ammonium hydroxide. The gradient was set as follows: 1 % B to 30 % B in 50 min, 45 % B in 10 min and 70 % B in 5 min. Fractions were collected at minute intervals and reduced using a vacuum concentrator. Dried peptides were reconstituted in 2 % MeCN, 0.1 % TFA and pooled to a total number of 12 fractions using a concatenation strategy covering the whole gradient^59^. An aliquot of each fraction (5-10 μg) was used for analysis of the global proteome and residual samples were dried in a vacuum concentrator prior to TiO_2_ phosphopeptide enrichment.

#### Phosphopeptide enrichment

Phosphopeptide enrichment was performed using a modified TiO_2_ batch protocol. In short, titanium dioxide beads (5 μm; GL Sciences, Japan) were sequentially washed with 120 μl 50% methanol, 300 μl ddH_2_O and 2x 300 μl binding solvent (1M glycolic acid, 70 % MeCN, 3 % TFA). In between, beads were spun down and the supernatant was discarded.

Dried peptides of each of the 12 fractions were individually resuspended in 150 μl binding solvent and incubated with the titanium dioxide beads at a bead to peptide ratio of 1:4 for 30 min at RT under continuous rotation. Bead-bound peptides were washed twice with binding solvent, 2x with washing solvent A (70 % MeCN, 3 % TFA) and 2x with washing solvent B (1 % MeCN, 0.1 % TFA). Phosphopeptides were eluted from the beads with 2x 150μl 0.3M NH_4_OH. The eluates were acidified by addition of TFA to a final concentration of 2 *%* and desalted using C18 StageTips^60^.

#### Liquid chromatography and tandem mass spectrometry

Global proteome and the phosphopeptide fractions were separated on an Ultimate 3000 RSLC nano-flow chromatography system using a pre-column for sample loading (PepMapAcclaim C18, 2 cm × 0.1 mm, 5 μm,) and a C18 analytical column (PepMapAcclaim C18, 50 cm × 0.75 mm, 2 μm, all Dionex, Thermo Fisher Scientific), applying a linear gradient over for 2 hours from 2 to 35% solvent B (80% acetonitrile, 0.1% formic acid; solvent A 0.1% formic acid) at a flow rate of 230 nl/min. Eluting peptides were analysed on an Orbitrap Fusion Lumos mass spectrometer equipped with EASY-Spray™ source (all Thermo Fisher Scientific), operated in a data-dependent acquisition mode with a cycle time of 3 s. FTMS1 spectra were recorded at a resolution of 120k, with an automated gain control (AGC) target of 200.000, and a max injection time of 50 ms. Precursors were filtered according to charge state (included charge states 2-6 z), and monoistopic peak assignment. Selected precursors were excluded from repeated fragmentation using a dynamic window (40 s, ± 10 ppm). The MS2 precursor were isolated with a quadrupole mass filter width of 1.2 m/z. For FTMS2, the Orbitrap was operated at 50k resolution, with an AGC target of 100.000 and a maximal injection time of 150 ms for global proteome samples ad 250 ms for phosphopeptide samples. Precursors were fragmented by high-energy collision dissociation (HCD) at a normalized collision energy (NCE) of 42 *%*.

### Transcriptomics analysis

#### Library preparation and sequencing

The amount of total RNA was quantified using the Qubit Fluorometric Quantitation system (Life Technologies) and the RNA integrity number (RIN) was determined using the Experion Automated Electrophoresis System (Bio-Rad).

For the co-culture experiments the isolated RNA (1μg) was processed using the SENSE mRNA-Seq Library Prep Kit V2 (Lexogen; #SKU 001.96) according to the manufacturer’s protocol. The libraries were sequenced on an Illumina HiSeq 2500 platform with 50 bp single-end reads to obtain on average 25 million reads per sample.

For the samples of the anti-CD20 study, RNA-seq libraries were prepared with the TruSeq Stranded mRNA LT sample preparation kit (Illumina) using both, Sciclone and Zephyr liquid handling robotics (PerkinElmer). Library concentrations were quantified with the Qubit Fluorometric Quantitation system (Life Technologies) and the size distribution was assessed using the Experion Automated Electrophoresis System (Bio-Rad). For sequencing, samples were diluted and pooled into NGS libraries in equimolar amounts. Expression profiling libraries were sequenced on Illumina HiSeq 3000/4000 instruments in 50-base-pair-single-end mode.

Base calls provided by the Illumina Real-Time Analysis (RTA) software were subsequently converted into BAM format (Illumina2bam) before de-multiplexing (BamIndexDecoder) into individual, sample-specific BAM files via Illumina2bam tools (1.17.3 https://github.com/wtsi-npg/illumina2bam).

### Data analysis and statistical information

All statistical tests were performed using R version 3.5.1^61^. The complete R scripts including the processed input data for all datasets and plots shown in this manuscript are available as Jupyter notebooks in Supplementary File 1.

#### Proteomics

RAW files were converted to the MGF files using ProteoWizard’s msConvert tool (version 3.0.9393) using vendor libraries^62^. Peak list files were searched using MSGF+^63^ (version 10089) and X!Tandem^64^ (version 2017.2.1.2). The precursor tolerance was set to 10 ppm, fragment tolerance to 0.01 Da for X!Tandem and machine type to QExactive with HCD fragmentation in MSGF+, and 1 missed cleavage was allowed. TMT tags and Carbamidomethylation of C were set as fixed modifications, Oxidation of M and Deamidation of N and Q, and Phosphorylation of S, T, and Y (TiO2 enriched samples only) as variable modifications. Searches were performed against the human SwissProt database (version 17-02), combined with sequences of common contaminants and reversed decoy sequences. Search results were filtered based on the target-decoy strategy at 0.01 FDR if both search engines identified the same sequences, diverging results were discarded, and spectra only identified by one search engines filtered at 0.001 FDR (search engine specific).

For quantitation all spectra with >30% interference based on total ion current were discarded. Isotope impurity correction and normalisation of intensity values was performed using the R Bioconductor package isobar^65^ (version 1.26). Protein abundance was estimated using the R Bioconductor package MSnbase^66^ (version 2.6.3) using the iPQF method and only peptides uniquely mapping to a single protein. Differential expression analysis was performed using the R Bioconductor package limma^67^ (version 3.35.3). The linear model included the MS run, patient, cell type (PBMCB vs. TAB) as factors next to the treatment group.

#### Transcriptomics

Bam files were converted to FASTQ format using the samtools package (https://github.com/samtools/samtools. version 0.1.19). The first 9 amino acids were removed using the fastx_trimmer tool (http://hannonlab.cshl.edu/fastx_toolkit, 0.0.14). Alignment was performed using the STAR aligner^68^ (version 2.5.3a, 23104886) with the output set to read counts. Alignment was performed against the Ensembl human genome 38 (version 88). Differential expression was assessed using the Bioconductor R package edgeR^69^ (version 3.22). Genes with less than 100 transcripts found in all samples were discarded prior to analysis. The linear model included the patient and cell type next to the treatment.

#### Pathway analysis

Pathway analysis was performed using the Camera algorithm^70^ as implemented in limma against the Molecular Signatures Database^71^ (MSIGDB, v6.1) hallmark^72^ and C2 canonical pathways collection. The analysis was performed separately for proteomics and transcriptomics data and merged using the Cytoscape^73^ Enrichment Map plugin^74^ (version 3.1.0).

Cell type abundance in whole tissue samples was estimated using xCell^34^ (R package version 1.1.0). Abundance of custom signatures was estimated using the ssGSEA approach^34,75^ as implemented in the Bioconductor R package GSVA^76^ (version 1.28.0).

#### Public datasets

To create our curated, key functional signatures we extracted highly correlating genes from the data sets from cutaneous melanoma samples of The Cancer Genome Atlas (TCGA)^77^ using the Bioconductor R Package geneRecommender (version 1.52). TCGA mRNA expression data were extracted using cBioPortal’s R interface using the CGDS-R package^78^ (version 1.2.5). Survival analysis was performed using the survival package (version 2.42-6, https://github.com/therneau/survival).

We used the data from Riaz *et al*. to evaluate the effects of anti-PD1 therapy on our signatures^39^. The TPM normalized expression values were downloaded from GEO (GSE91061). Signature expression levels were estimated using the ssGSEA approach and cell type abundances estimated using xCell (see above).

scRNA-seq from Sade-Feldman *et al*.^29^ was directly downloaded from GEO as TPM normalized expression values (GSE120575). The complete data analysis was performed in R using Seurat 2.3.4^79^. The data was normalized using the “LogNormalize” function with a scale factor of 10,000. Variable genes were detected using the “LogVMR” dispersion function and the “ExpMean” function to calculate means. The x.low.cutoff was set to 0.0125, x.high.cutoff to 3, and the y.cutoff to 0.5. Next, the data was scaled regressing out the patient identifier. Out of 100 calculated principal components, the first 35 were used for further analysis. Cells were clustered based on the principal components using a resolution of 0.6. Subsequent cell types were annotated based on canonical markers and all B cell-like cells retained. B cells were again clustered using a resolution of 0.9. Clusters were annotated based on the found markers (function FindMarkers, min.pct set to 0.25), as well as the expression of canonical B cell markers. Signature expression levels were estimated using the ssGSEA function (see above) based on the average gene expressions. The complete workflow can be found in Supplementary File 1 as a Jupyter notebook.

### Histology

Tissue arrays consisted of human melanoma metastases from cutaneous and subcutaneous sites with 0.5-mm cores. All tumor samples were obtained with informed patients’ consent and retrieved from the pathology files of the Medical University of Vienna (ethics vote: 405/2006). Representative tumor areas were selected based on the review of hematoxylin and eosin-stained sections from formalin-fixed paraffin-embedded (FFPE) blocks by two authors of this study (PP, SNW), respectively. In selected tumor samples, S-100, HMB45 or triple CSPG/β3 integrin/HMB45 melanoma marker immunostainings were done to identify tumor cells and to confirm the diagnosis. Criteria for exclusion of cores were: missing, necrotic or extensive hemorrhagic tumor areas and insufficient multiplex staining. Using these criteria, of the 216 cores stained, 155 cores representing 58 melanomas from 30 different patients were included in the final analysis.

#### Multiplex IHC staining

4 μm sections from full FFPE blocks that were used to generate the TMA or from control tonsil tissue were utilized for both, the initial establishment of staining conditions for each individual primary antibody (Ab) and the successive optimization of multiplex staining. In a first step, primary Abs against the following antigens were established as single stains initially on human tonsil tissue and thereafter on metastatic human melanoma tumors: CD19, CD20, CD5, CD27, CD38, CD138, CD8, FoxP3 and IL10 (Supplementary Table 5).

In a second step, multiplex immunostains were established essentially as described^80,81^. Random integration of sequential Abs within a multiplex panel may lead to imbalanced signals, incomplete staining through interference with previously applied tyramide signal amplification (TSA), disruption of epitopes, and removal of TSA fluorophores because of repetitive antigen-retrievals at high temperature^80,81^. Therefore, each Ab was tested individually for its optimal position in the sequence of multiplex staining to minimize interference with previous Ab-TSA complexes or by alteration of epitopes.

For multiplex immunostains, 4 μm sections were deparaffinized and antigen retrieval was performed in heated citrate buffer (pH 6.0) and/or Tris-EDTA buffer (pH 9) for 30 min. Thereafter, sections were fixed with 7.5% neutralized formaldehyde (SAV Liquid Production GmbH). Each section was subjected to 6 successive rounds of Ab staining, each consisting of protein blocking with 20% normal goat serum (Dako) in PBS, incubation with primary Abs, biotinylated anti-mouse/rabbit secondary antibodies and Streptavidin-HRP (Dako), followed by TSA visualization with fluorophores Opal 520, Opal 540, Opal 570, Opal 620, Opal 650, and Opal 690 (PerkinElmer) diluted in 1X Plus Amplification Diluent (PerkinElmer), Ab-TSA complex-“stripping” in heated citrate buffer (pH 6.0) and/or Tris-EDTA buffer (pH 9) for 30 min and fixation with 7.5% neutralized formaldehyde. Thereafter, nuclei were counterstained with DAPI (PerkinElmer) and sections mounted with PermaFluor fluorescence mounting medium (Thermo Fisher Scientific).

Respective stainings without primary antibodies were used as a negative control. Along with tissue arrays, serial sections of melanoma specimens and normal controls were stained to assess reproducibility. At equal Ab-concentrations, TSA-based visualization is expected to yield a higher number of positive cells as compared to conventional immunofluorescence. We therefore established TSA-based visualization of primary Abs on control tonsil tissue, the golden standard for lymphocyte antigen detection in pathology, and performed a comparison for each Ab to validated staining patterns in human tonsil (as to the Human Protein Atlas^82^, available from www.poroteinatlas.org). Thereafter, we balanced the signal through dilution of the primary Abs to obtain staining levels and cell frequencies comparable to conventional immunofluorescence staining. The dilution of the CD19 antibody was optimized to allowing for the detection of CD19^low^ plasma cell-like cells as compared to patterns and frequencies obtained by CD138+ pooled lgA/lgG+ stainings on human tonsil and melanoma^17,19^. In multiplex stainings, single primary Ab stainings were run in parallel to control for false positive results through incomplete Ab-TSA complex-“stripping” and false negative results through antigen masking (by incubation with multiple primary Abs, “umbrella-effect”). Spillover effects were controlled for anti-CD20-Ab stainings on tonsil with different Opal fluorophores by signal detection in adjacent components/channels and thereafter for exposure time settings upon acquisition of multiplex-stained tissue sections.

#### Tissue imaging, spectral unmixing and phenotyping

Multiplexed slides were scanned on a Vectra Multispectral Imaging System version 2 (Perkin Elmer) as described^80,81^. Briefly, a spectral library from spectral peaks emitted by each fluorophore from single stained slides was created with the inform Advanced Image Analysis software (InForm 2.4, PerkinElmer) and used for spectral unmixing of multispectral images allowing for identification of all markers of interest. Autofluorescence was determined on an unstained representative study sample. To subtype TAB in TMA, cells were phenotyped as (i) CD19^+^ CD20^-^ CD38^+^ CD138^-^ plasmablast-like, CD19^+^ CD20^-^ CD138^+^ plasma cell-like, CD19^+^ CD20^+^ CD38^-^ CD138^-^ memory B cell-like, CD20^+^ CD38^+^ CD138^-^ CD5^-^ germinal center B cell-like, CD19^+^ CD20^-^ CD38^-^ CD138^-^ CD27^+^ activated B cell-like, CD20^+^ CD19^-^ CD138^-^ CD5^+^ transitional cell-like TAB and (ii) other cells. The staining protocol has been optimized for detection of CD19. Though CD19^low^ plasma cell-like cells could be detected at significant numbers, they may still be underrepresented in our staining data. The same may also be true for the detection of activated B cell-like cells, as expression of CD27 has been reported to be downregulated on TAB^43,83^. Human melanoma cells can express CD38^84^ and we observed some cores with slight CD38 immunoreactivity of melanoma cells. In these cores tumor cell areas were excluded with the “exclusion area tool” of InForm. After adaptive cell segmentation, a selection of around 25 representative original multispectral images was used to set cut-off values for each fluorophore/antibody staining. All phenotyping and subsequent quantifications were performed blinded to the sample identity.

## References

1. Ascierto, P. A. et al. Survival Outcomes in Patients With Previously Untreated BRAF Wild-Type Advanced Melanoma Treated With Nivolumab Therapy: Three-Year Follow-up of a Randomized Phase 3 Trial. JAMA Oncol (2018). doi:10.1001/jamaoncol.2018.4514

2. de Visser, K. E., Korets, L. V. & Coussens, L. M. De novo carcinogenesis promoted by chronic inflammation is B lymphocyte dependent. Cancer Cell 7, 411–423 (2005).

3. Ammirante, M., Luo, J.-L., Grivennikov, S., Nedospasov, S. & Karin, M. B-cell-derived lymphotoxin promotes castration-resistant prostate cancer. Nature 464, 302–305 (2010).

4. Shalapour, S. et al. Immunosuppressive plasma cells impede T-cell-dependent immunogenic chemotherapy. Nature 521, 94–98 (2015).

5. Affara, N. I. et al. B cells regulate macrophage phenotype and response to chemotherapy in squamous carcinomas. Cancer Cell 25, 809–821 (2014).

6. Gunderson, A. J. et al. Bruton Tyrosine Kinase-Dependent Immune Cell Cross-talk Drives Pancreas Cancer. Cancer Discov. 6, 270–285 (2016).

7. Shalapour, S. et al. Inflammation-induced IgA+ cells dismantle anti-liver cancer immunity. Nature 551, 340–345 (2017).

8. Mauri, C. & Menon, M. Human regulatory B cells in health and disease: therapeutic potential. J. Clin. Invest. 127, 772–779 (2017).

9. Shimabukuro-Vornhagen, A. et al. Characterization of tumor-associated B-cell subsets in patients with colorectal cancer. Oncotarget 5, 4651–4664 (2014).

10. Zhou, X., Su, Y.-X., Lao, X.-M., Liang, Y.-J. & Liao, G.-Q. CD19(+)IL-10(+) regulatory B cells affect survival of tongue squamous cell carcinoma patients and induce resting CD4(+) T cells to CD4(+)Foxp3(+) regulatory T cells. Oral Oncol. 53, 27–35 (2016).

11. Lindner, S. et al. Interleukin 21-induced granzyme B-expressing B cells infiltrate tumors and regulate T cells. Cancer Res. 73, 2468–2479 (2013).

12. Wei, X. et al. Regulatory B cells contribute to the impaired antitumor immunity in ovarian cancer patients. Tumour Biol. 37, 6581–6588 (2016).

13. Xiao, X. et al. PD-1hi Identifies a Novel Regulatory B-cell Population in Human Hepatoma That Promotes Disease Progression. Cancer Discov. 6, 546–559 (2016).

14. Liu, J. et al. Aberrant frequency of IL-10-producing B cells and its association with Treg and MDSC cells in Non Small Cell Lung Carcinoma patients. Hum. Immunol. 77, 84–89 (2016).

15. Ishigami, E. et al. Coexistence of regulatory B cells and regulatory T cells in tumor-infiltrating lymphocyte aggregates is a prognostic factor in patients with breast cancer. Breast Cancer (2018). doi:10.1007/s12282-018-0910-4

16. Somasundaram, R. et al. Tumor-associated B-cells induce tumor heterogeneity and therapy resistance. Nat. Commun. 8, 607 (2017).

17. Erdag, G. et al. Immunotype and immunohistologic characteristics of tumor-infiltrating immune cells are associated with clinical outcome in metastatic melanoma. Cancer Res. 72, 1070–1080 (2012).

18. Ladányi, A. Prognostic and predictive significance of immune cells infiltrating cutaneous melanoma. Pigment Cell Melanoma Res. 28, 490–500 (2015).

19. Bosisio, F. M. et al. Plasma cells in primary melanoma. Prognostic significance and possible role of IgA. Mod. Pathol. 29, 347–358 (2016).

20. Winkler, J. K., Schiller, M., Bender, C., Enk, A. H. & Hassel, J. C. Rituximab as a therapeutic option for patients with advanced melanoma. Cancer Immunol. Immunother. 67, 917–924 (2018).

21. Rosser, E. C. et al. Regulatory B cells are induced by gut microbiota-driven interleukin-1β and interleukin-6 production. Nat. Med. 20, 1334–1339 (2014).

22. Wang, R.-X. et al. Interleukin-35 induces regulatory B cells that suppress autoimmune disease. Nat. Med. 20, 633–641 (2014).

23. Rosser, E. C. & Mauri, C. Regulatory B cells: origin, phenotype, and function. Immunity 42, 607–612 (2015).

24. Sumimoto, H., Imabayashi, F., Iwata, T. & Kawakami, Y. The BRAF-MAPK signaling pathway is essential for cancer-immune evasion in human melanoma cells. J. Exp. Med. 203, 1651–1656 (2006).

25. Wang, Z. et al. Tumor-derived IL-35 promotes tumor growth by enhancing myeloid cell accumulation and angiogenesis. J. Immunol. 190, 2415–2423 (2013).

26. Bindea, G. et al. Spatiotemporal dynamics of intratumoral immune cells reveal the immune landscape in human cancer. Immunity 39, 782–795 (2013).

27. Hoesel, B. & Schmid, J. A. The complexity of NF-κB signaling in inflammation and cancer. Mol. Cancer 12, 86 (2013).

28. Wu, G. & Haw, R. Functional Interaction Network Construction and Analysis for Disease Discovery. Methods Mol. Biol. 1558, 235–253 (2017).

29. Sade-Feldman, M. et al. Defining T Cell States Associated with Response to Checkpoint Immunotherapy in Melanoma. Cell 175, 998–1013.e20 (2018).

30. de Oliveira, C. E. C. et al. CC chemokine receptor 5: the interface of host immunity and cancer. Dis. Markers 2014, 126954 (2014).

31. Bystry, R. S., Aluvihare, V., Welch, K. A., Kallikourdis, M. & Betz, A. G. B cells and professional APCs recruit regulatory T cells via CCL4. Nat. Immunol. 2, 1126–1132 (2001).

32. Facciabene, A. et al. Tumour hypoxia promotes tolerance and angiogenesis via CCL28 and T(reg) cells. Nature 475, 226–230 (2011).

33. Günther, C., Carballido-Perrig, N., Kaesler, S., Carballido, J. M. & Biedermann, T. CXCL16 and CXCR6 are upregulated in psoriasis and mediate cutaneous recruitment of human CD8+ T cells. J. Invest. Dermatol. 132, 626–634 (2012).

34. Aran, D., Hu, Z. & Butte, A. J. xCell: digitally portraying the tissue cellular heterogeneity landscape. Genome Biol. 18, 220 (2017).

35. Danaher, P. et al. Pan-cancer adaptive immune resistance as defined by the Tumor Inflammation Signature (TIS): results from The Cancer Genome Atlas (TCGA). J Immunother Cancer 6, 63 (2018).

36. Ayers, M. et al. IFN-γ-related mRNA profile predicts clinical response to PD-1 blockade. J. Clin. Invest. 127, 2930–2940 (2017).

37. Elia, A. R., Caputo, S. & Bellone, M. Immune Checkpoint-Mediated Interactions Between Cancer and Immune Cells in Prostate Adenocarcinoma and Melanoma. Front. Immunol. 9, 1786 (2018).

38. Bolen, C. R. et al. Mutation load and an effector T-cell gene signature may distinguish immunologically distinct and clinically relevant lymphoma subsets. Blood Adv 1, 1884–1890 (2017).

39. Riaz, N. et al. Tumor and Microenvironment Evolution during Immunotherapy with Nivolumab. Cell 171, 934–949.e16 (2017).

40. Pinc, A. et al. Targeting CD20 in melanoma patients at high risk of disease recurrence. Mol. Ther. 20, 1056–1062 (2012).

41. Chen, D. S. & Mellman, I. Elements of cancer immunity and the cancer-immune set point. Nature 541, 321–330 (2017).

42. Hugo, W. et al. Non-genomic and Immune Evolution of Melanoma Acquiring MAPKi Resistance. Cell 162, 1271–1285 (2015).

43. Hegde, P. S., Karanikas, V. & Evers, S. The Where, the When, and the How of Immune Monitoring for Cancer Immunotherapies in the Era of Checkpoint Inhibition. Clin. Cancer Res. 22, 1865–1874 (2016).

44. Li, R., Patterson, K. R. & Bar-Or, A. Reassessing B cell contributions in multiple sclerosis. Nat. Immunol. 19, 696–707 (2018).

45. Kessel, A. et al. Human CD19(+)CD25(high) B regulatory cells suppress proliferation of CD4(+) T cells and enhance Foxp3 and CTLA-4 expression in T-regulatory cells. Autoimmun. Rev. 11, 670–677 (2012).

46. Parekh, V. V. et al. B Cells Activated by Lipopolysaccharide, But Not By Anti-Ig and Anti-CD40 Antibody, Induce Anergy in CD8+ T Cells: Role of TGF- 1. The Journal of Immunology 170, 5897–5911 (2003).

47. Matsumoto, M. et al. Interleukin-10-producing plasmablasts exert regulatory function in autoimmune inflammation. Immunity 41, 1040–1051 (2014).

48. Sun, C.-M., Deriaud, E., Leclerc, C. & Lo-Man, R. Upon TLR9 signaling, CD5+ B cells control the IL-12-dependent Th1-priming capacity of neonatal DCs. Immunity 22, 467–477 (2005).

49. Iwata, Y. et al. Characterization of a rare IL-10-competent B-cell subset in humans that parallels mouse regulatory B10 cells. Blood 117, 530–541 (2011).

50. Biswas, S. K. & Mantovani, A. Macrophage plasticity and interaction with lymphocyte subsets: cancer as a paradigm. Nat. Immunol. 11, 889–896 (2010).

51. Mao, H. et al. CD19CD27 Plasmablasts Suppress Harmful Th17 Inflammation Through Interleukin 10 Pathway in Colorectal Cancer. DNA Cell Biol. 36, 870–877 (2017).

52. Zhu, Y., Yao, S. & Chen, L. Cell surface signaling molecules in the control of immune responses: a tide model. Immunity 34, 466–478 (2011).

53. Kolesnikov, N. et al. ArrayExpress update--simplifying data submissions. Nucleic Acids Res. 43, D1113–6 (2015).

54. Perez-Riverol, Y. et al. The PRIDE database and related tools and resources in 2019: improving support for quantification data. Nucleic Acids Res. (2018). doi:10.1093/nar/gky1106

55. Maurer, M. et al. Comprehensive comparative and semiquantitative proteome of a very low number of native and matched epstein-barr-virus-transformed B lymphocytes infiltrating human melanoma. J. Proteome Res. 13, 2830–2845 (2014).

56. Maurer, M. et al. Combining filter-aided sample preparation and pseudoshotgun technology to profile the proteome of a low number of early passage human melanoma cells. J. Proteome Res. 12, 1040–1048 (2013).

57. Fraussen, J. et al. A novel method for making human monoclonal antibodies. J. Autoimmun. 35, 130–134 (2010).

58. Wiśniewski, J. R., Zougman, A., Nagaraj, N. & Mann, M. Universal sample preparation method for proteome analysis. Nat. Methods 6, 359–362 (2009).

59. Batth, T. S., Francavilla, C. & Olsen, J. V. Off-line high-pH reversed-phase fractionation for in-depth phosphoproteomics. J. Proteome Res. 13, 6176–6186 (2014).

60. Rappsilber, J., Mann, M. & Ishihama, Y. Protocol for micro-purification, enrichment, pre-fractionation and storage of peptides for proteomics using StageTips. Nat. Protoc. 2, 1896–1906 (2007).

61. R Core Team. R: A Language and Environment for Statistical Computing. (2018).

62. Adusumilli, R. & Mallick, P. Data Conversion with ProteoWizard msConvert. Methods Mol. Biol. 1550, 339–368 (2017).

63. Kim, S. & Pevzner, P. A. MS-GF+ makes progress towards a universal database search tool for proteomics. Nat. Commun. 5, 5277 (2014).

64. MacLean, B., Eng, J. K., Beavis, R. C. & McIntosh, M. General framework for developing and evaluating database scoring algorithms using the TANDEM search engine. Bioinformatics 22, 2830–2832 (2006).

65. Breitwieser, F. P. et al. General statistical modeling of data from protein relative expression isobaric tags. J. Proteome Res. 10, 2758–2766 (2011).

66. Gatto, L. & Lilley, K. S. MSnbase-an R/Bioconductor package for isobaric tagged mass spectrometry data visualization, processing and quantitation. Bioinformatics 28, 288–289 (2012).

67. Ritchie, M. E. et al. limma powers differential expression analyses for RNA-sequencing and microarray studies. Nucleic Acids Res. 43, e47 (2015).

68. Dobin, A. et al. STAR: ultrafast universal RNA-seq aligner. Bioinformatics 29, 15–21 (2013).

69. McCarthy, D. J., Chen, Y. & Smyth, G. K. Differential expression analysis of multifactor RNA-Seq experiments with respect to biological variation. Nucleic Acids Res. 40, 4288–4297 (2012).

70. Wu, D. & Smyth, G. K. Camera: a competitive gene set test accounting for inter-gene correlation. Nucleic Acids Res. 40, e133 (2012).

71. Liberzon, A. et al. Molecular signatures database (MSigDB) 3.0. Bioinformatics 27, 1739–1740 (2011).

72. Liberzon, A. et al. The Molecular Signatures Database (MSigDB) hallmark gene set collection. Cell Syst 1, 417–425 (2015).

73. Shannon, P. et al. Cytoscape: a software environment for integrated models of biomolecular interaction networks. Genome Res. 13, 2498–2504 (2003).

74. Merico, D., Isserlin, R., Stueker, O., Emili, A. & Bader, G. D. Enrichment map: a network-based method for gene-set enrichment visualization and interpretation. PLoS One 5, e13984 (2010).

75. Barbie, D. A. et al. Systematic RNA interference reveals that oncogenic KRAS-driven cancers require TBK1. Nature 462, 108–112 (2009).

76. Hänzelmann, S., Castelo, R. & Guinney, J. GSVA: gene set variation analysis for microarray and RNA-seq data. BMC Bioinformatics 14, 7 (2013).

77. Cancer Genome Atlas Network. Genomic Classification of Cutaneous Melanoma. Cell 161, 1681–1696 (2015).

78. Cerami, E. et al. The cBio cancer genomics portal: an open platform for exploring multidimensional cancer genomics data. Cancer Discov. 2, 401–404 (2012).

79. Butler, A., Hoffman, P., Smibert, P., Papalexi, E. & Satija, R. Integrating single-cell transcriptomic data across different conditions, technologies, and species. Nat. Biotechnol. 36, 411 (2018).

80. Carstens, J. L. et al. Spatial computation of intratumoral T cells correlates with survival of patients with pancreatic cancer. Nat. Commun. 8, 15095 (2017).

81. Gorris, M. A. J. et al. Eight-Color Multiplex Immunohistochemistry for Simultaneous Detection of Multiple Immune Checkpoint Molecules within the Tumor Microenvironment. J. Immunol. 200, 347–354 (2018).

82. Uhlén, M. et al. Proteomics. Tissue-based map of the human proteome. Science 347, 1260419 (2015).

83. Nielsen, J. S. et al. CD20+ tumor-infiltrating lymphocytes have an atypical CD27- memory phenotype and together with CD8+ T cells promote favorable prognosis in ovarian cancer. Clin. Cancer Res. 18, 3281–3292 (2012).

84. Morandi, F. et al. A non-canonical adenosinergic pathway led by CD38 in human melanoma cells induces suppression of T cell proliferation. Oncotarget 6, 25602–25618 (2015).

