## Supplementary Figures and Methods for "B cells sustain inflammation and predict response to immune checkpoint blockade in human melanoma"

### Table of Contents

**Table of Contents** 2

**Supplementary Figures** 3

Supplementary Figure 1 3

Supplementary Figure 2 4

Supplementary Figure 3 5

Supplementary Figure 4 6

Supplementary Figure 5 7

Supplementary Figure 6 8

Supplementary Figure 7 9

Supplementary Figure 8 10

**Supplementary Methods** 11

FACS Gating Strategy 11

### Supplementary Figures

#### Supplementary Figure 1

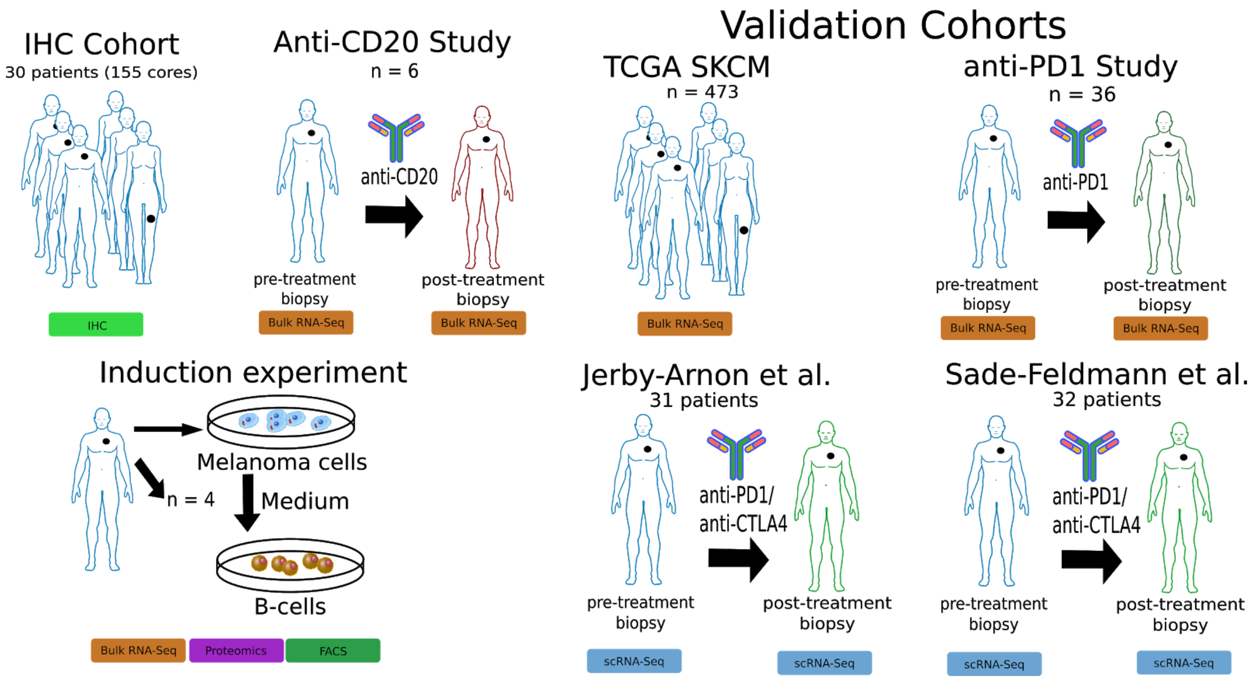


**Patient cohorts evaluated in this study.** Melanoma TAB phenotypes were characterised using TMAs from 30 patients with 155 cores from 58 different metastatic melanomas. Changes induced by melanoma cells in TAB were evaluated in vitro using autologous B cells from 4 patients and melanoma conditioned medium from one patient, all with metastatic melanoma. Finally, the loss of B cells was evaluated using clinical samples from an anti-CD20 study performed by our group. All findings were validated using 3 public datasets: the TCGA skin cutaneous melanoma cohort, whole tissue RNA-seq data from melanoma patients pre- and on-anti-PD1 therapy and scRNA-seq data from melanoma lesions from two independent studies.

#### Supplementary Figure 2

A)

| TAB subpopulations | IHC-staining pattern |
|---|---|
| Plasmablast-like | $CD19^{+}$ $CD20^{-}$ $CD38^{+}$ $CD138^{-}$ |
| Plasma cell-like | $CD19^{+}$ $CD20^{-}$ $CD138^{+}$ |
| Memory B cell-like | $CD19^{+}$ $CD20^{+}$ $CD38^{-}$ $CD138^{-}$ |
| Germinal center B cell-like | $CD20^{+}$ $CD38^{+}$ $CD138^{-}$ $CD5^{-}$ |
| Activated B cell-like | $CD19^{+}$ $CD20^{-}$ $CD38^{-}$ $CD138^{-}$ $CD27^{+}$ |
| Transitional cell-like | $CD20^{+}$ $CD19^{-}$ $CD138^{-}$ $CD5^{+}$ |

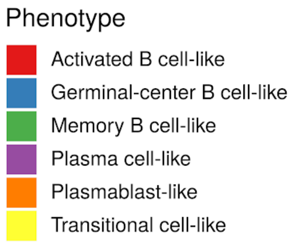


B)

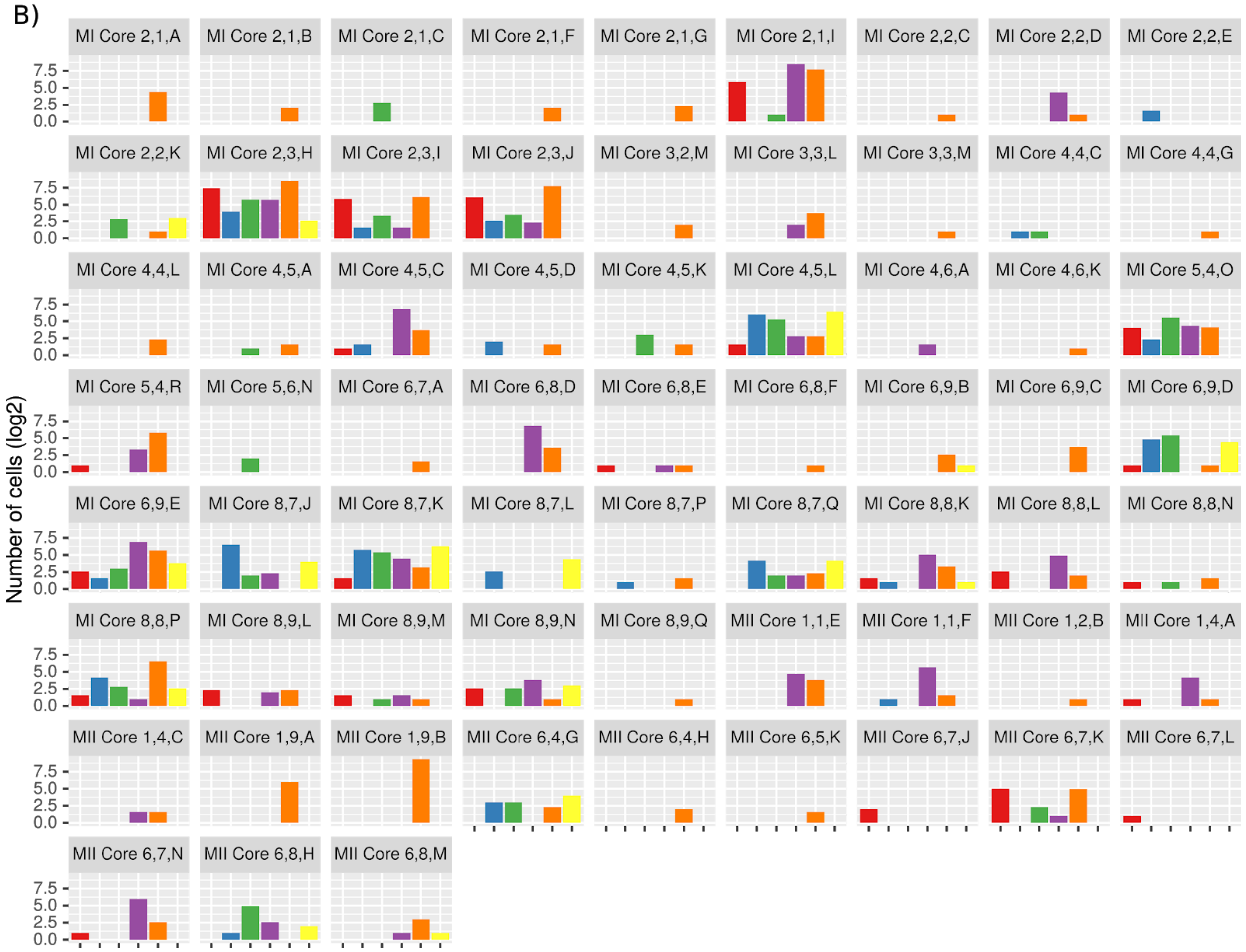


**Phenotypic characterization of melanoma TAB.** A) Marker combinations used to identify TAB subpopulations by multiplex immunostaining. B) Frequency of TAB subpopulations in the 155 metastatic melanoma cores. Only cores with at least one characterized cell are shown. Identified cells per core (log2-transformed) are shown for every core.

Supplementary Figure 3

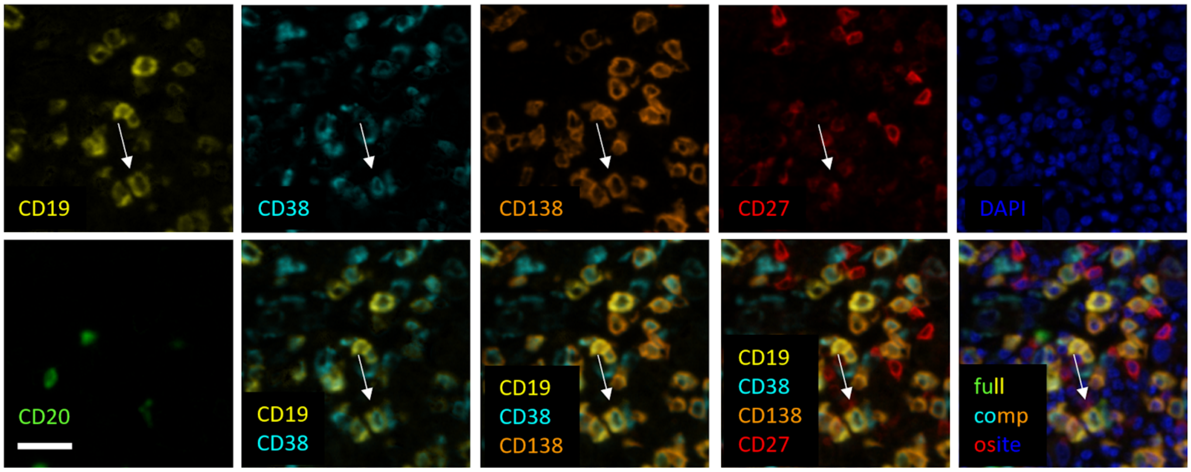


**Multiplex immunostaining to identify plasma cell-like TAB.** Identification of plasma cell-like TAB. Composite image together with DAPI nuclear staining (bottom right) and images for each of the individual markers and different combinations from the composite image. Arrows depict one of several plasma cell-like TAB. Scale bar represents 20µm.

### Supplementary Figure 4

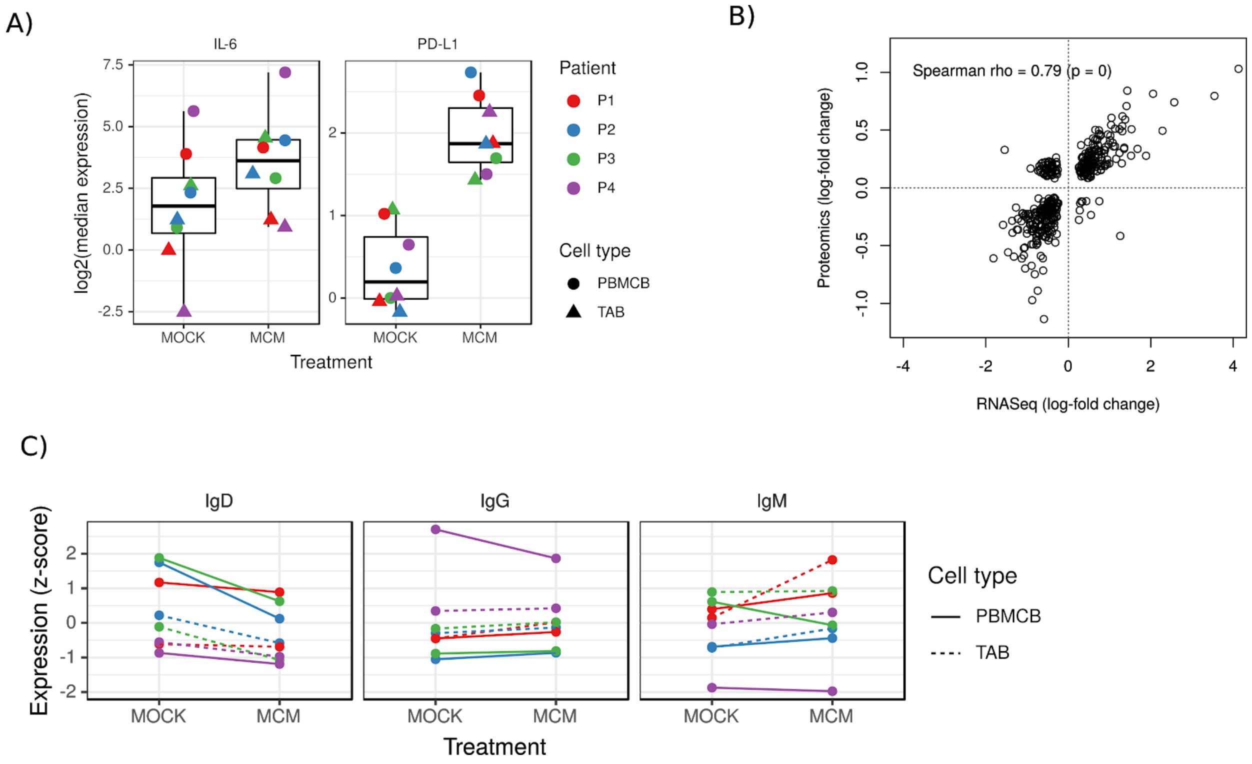


**Induction experiments, significantly regulated gene/proteins and FACS data in B cells induced with melanoma conditioned medium. A)** Induction of PD-L1 and IL-6 expression in peripheral blood B cells (PBMCB)- and tumour -derived B cells (TAB) by melanoma conditioned (MCM) and control (MOCK) medium. mRNA transcripts were determined by RT-qPCR with levels indicated as RQ values normalized to b-actin (Taq Man gene expression assays for IL-6: Hs00174131_m1, PD-L1: Hs00204257_m1, all ThermoFisher). Lower and upper hinges of the boxplots represent the first and third quartiles. Whiskers represent the highest and lowest value no further than 1.5 times the interquartile range of the first and third quartile. Colored points represent the actually observed values. Regulation through MCM was significant for both genes (paired Wilcoxon signed rank test, Bonferroni adjusted p = 0.016 for both). **B)** Fold changes (log2 transformed) of genes/proteins identified as significantly regulated showed a high correlation between proteomics and RNA-seq results. One group of proteins was consistently upregulated while the corresponding transcripts were downregulated (top-left corner). These all belong to proteins associated with metabolic pathways. **C)** Additional FACS-estimated expression levels of IgD, IgG, and IgM on peripheral blood B cells (PBMCB)- and tumour -derived B cells (TAB) by melanoma conditioned (MCM) and control (MOCK) medium.

#### Supplementary Figure 5

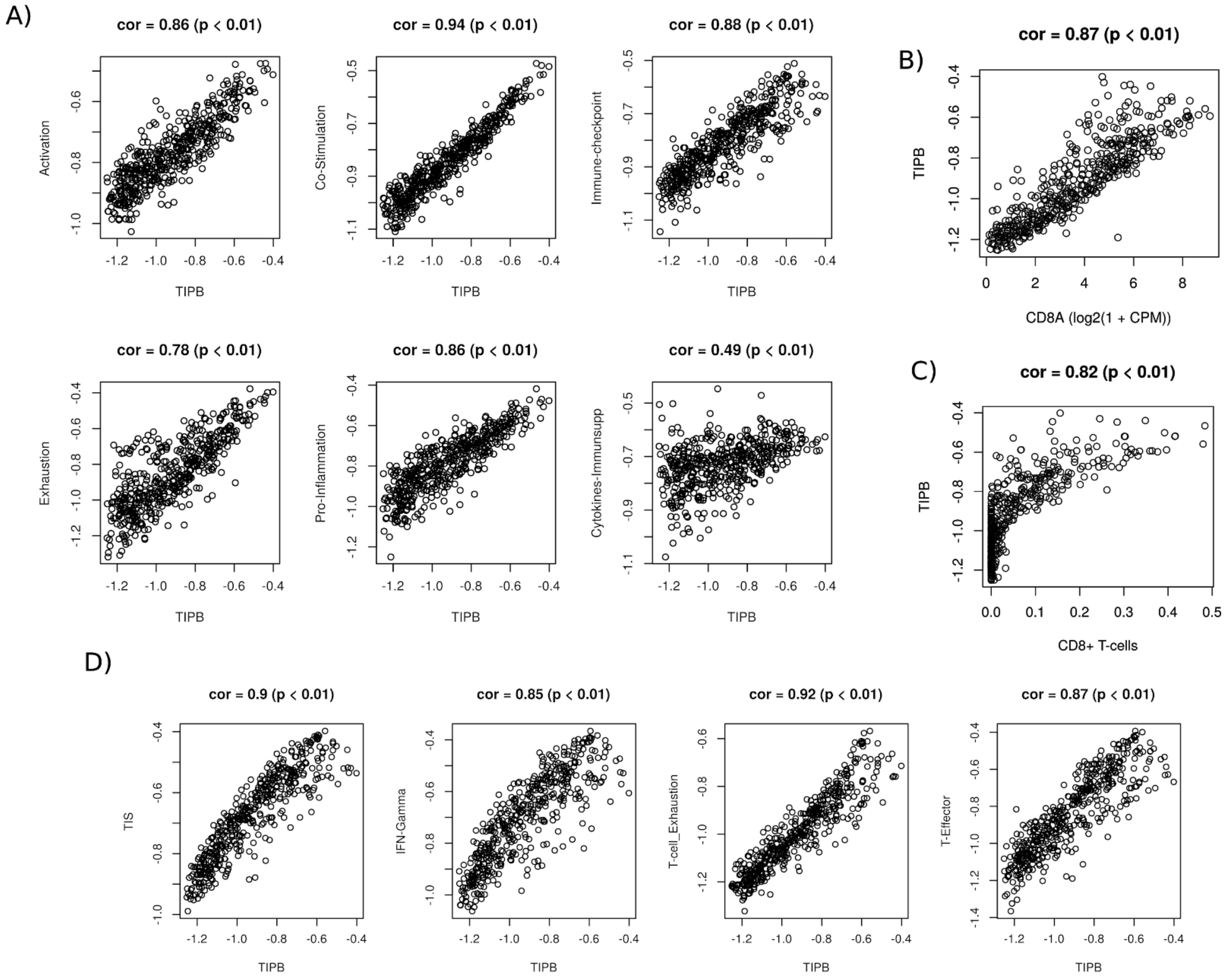


**Validation of the TIPB and the functional signatures in the TCGA skin cutaneous melanoma cohort. A)** Correlation of the functional signatures with the TIPB signature. **B)** Correlation of the TIPB signature with the expression of CD8A and **C)** with the xCell estimated abundance of CD8+ T-cells. **D)** Correlation of the TIPB signature with established signatures describing the inflammation in the TME and T cell function and phenotype (tumor inflammatory score (TIS), interferon (IFN) gamma, T cell exhaustion, T cell effector (T-effector)) highly correlated with our TIPB signature. All correlation coefficients and p-values refer to the Spearman correlation coefficient.

#### Supplementary Figure 6

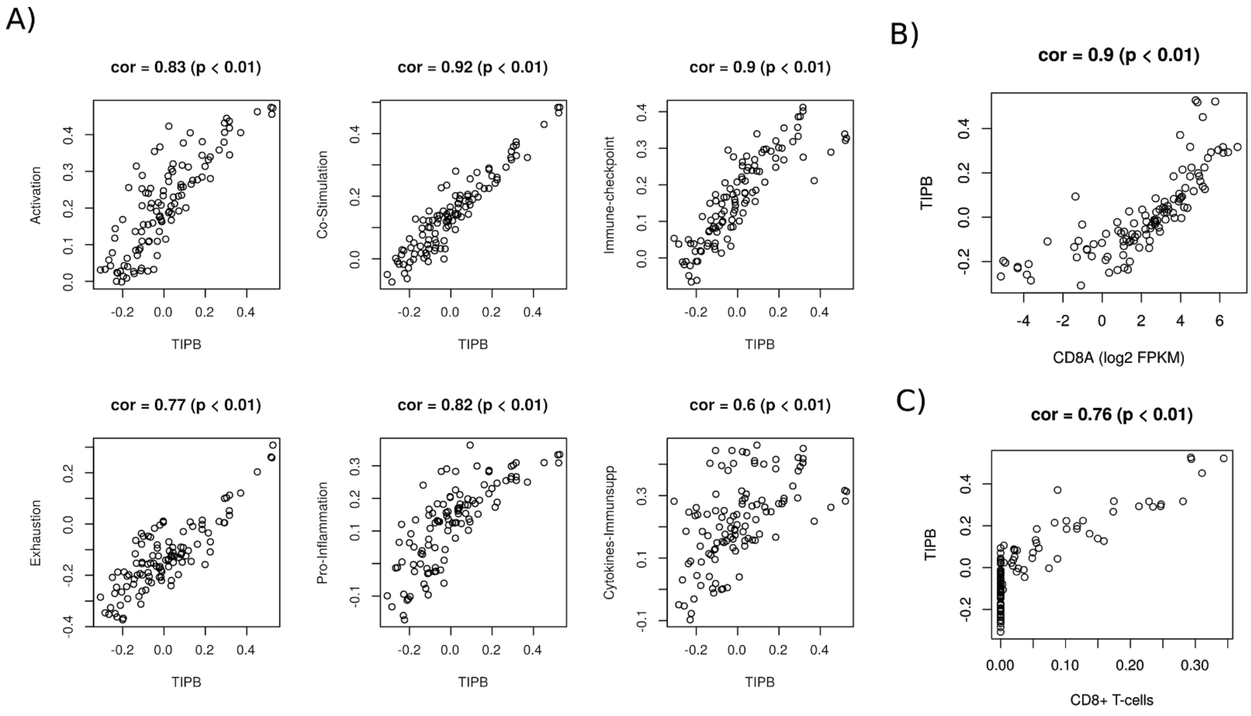


**Validation of the TIPB and the functional signatures in a melanoma cohort treated with anti-PD1 (Riaz et al. dataset). A)** Correlation of functional signatures as estimated by ssGSEA with the TIPB signature. **B)** Correlation of the TIPB signature with the expression of CD8A and **C)** the xCell estimated abundance of CD8+ T cells.

#### Supplementary Figure 7

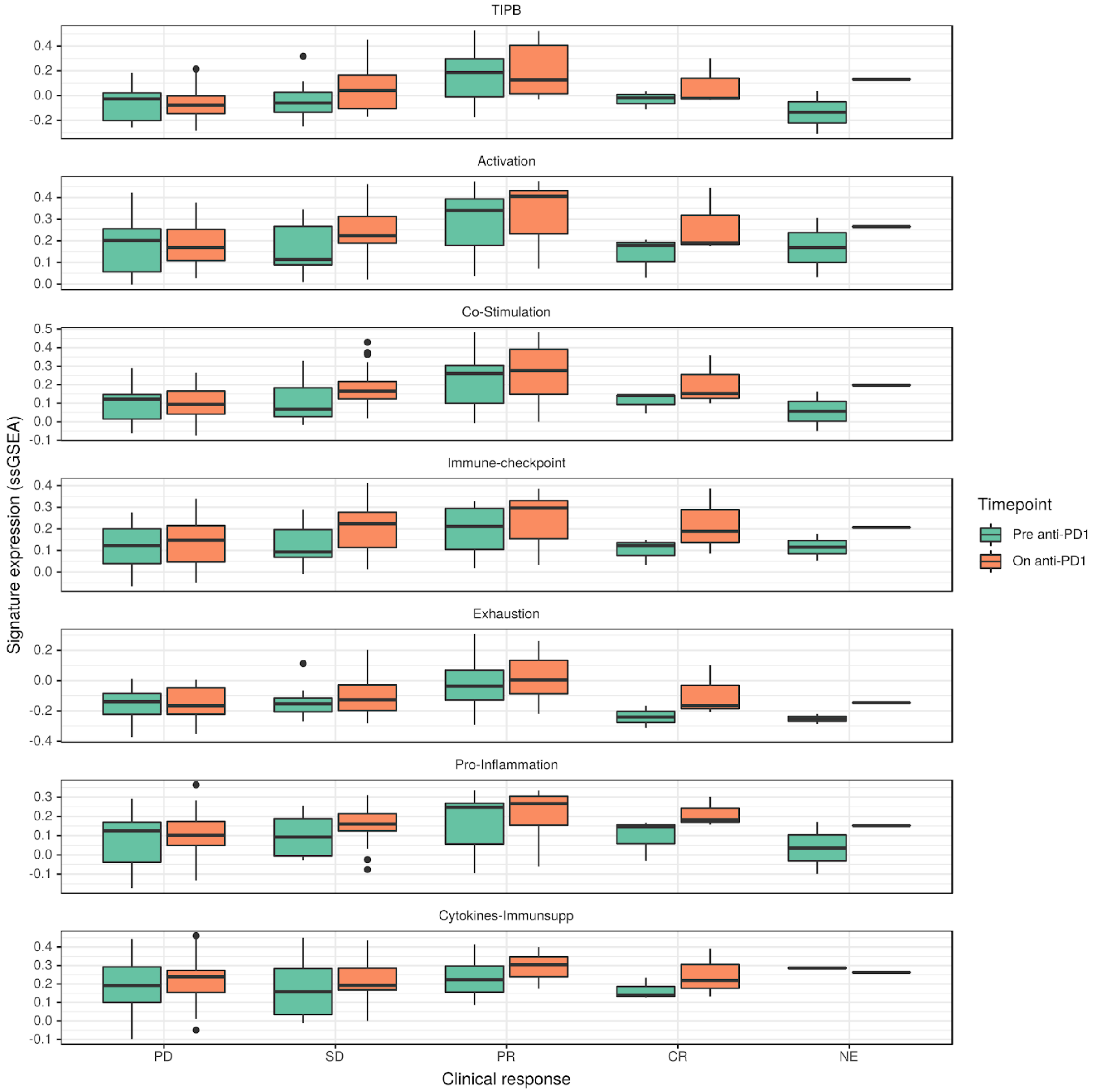


**Expression of the TIPB signature and the functional signatures before and on anti-PD1 therapy versus clinical response (Riaz *et al.* dataset).** (PD = progressive disease, SD = stable disease, PR = partial response, CR = complete response, NE = not evaluated)

#### Supplementary Figure 8

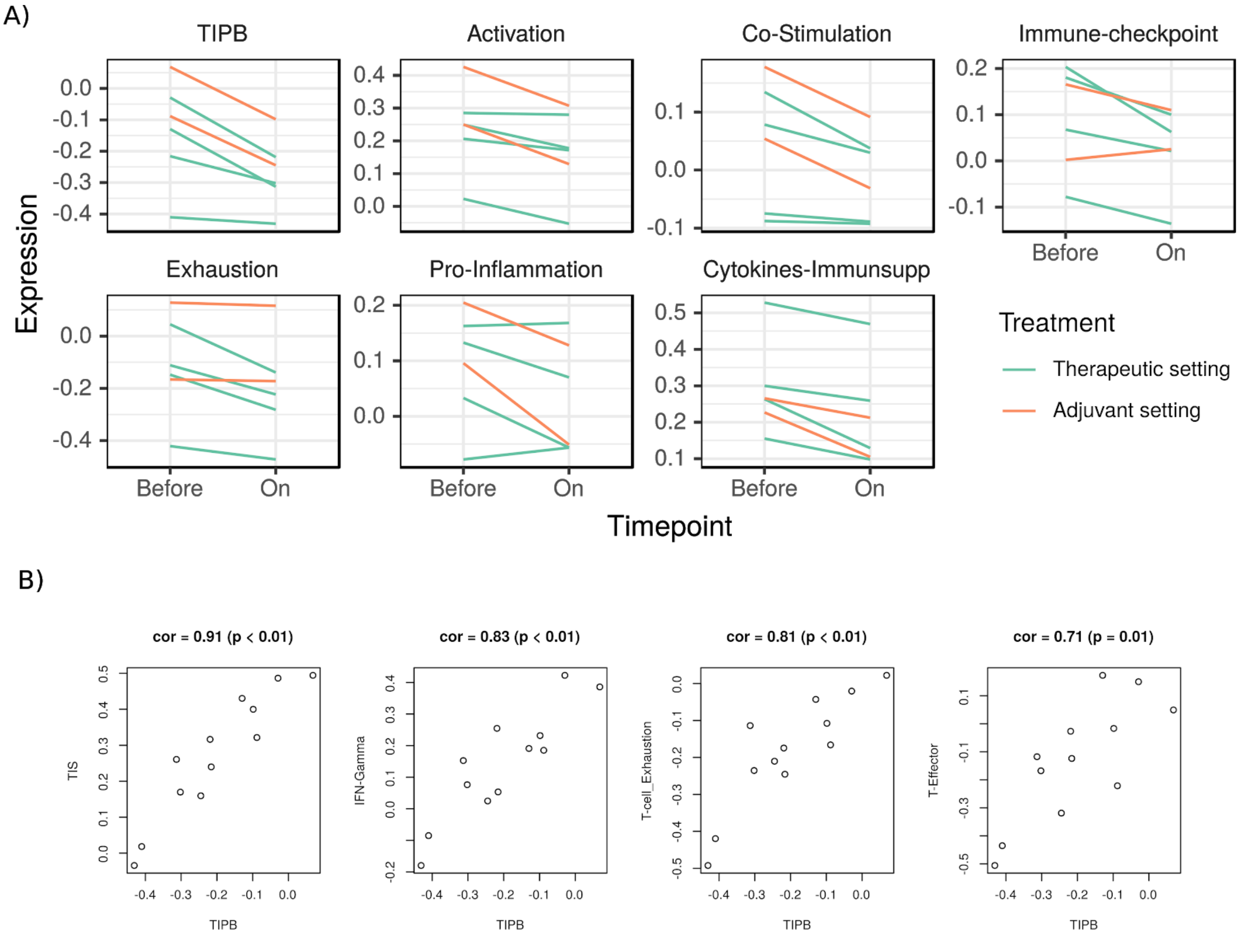


**Validation of the TIPB and the functional signatures in the anti-CD20 clinical study samples.** A) Estimated abundance (ssGSEA) of the TIPB signature and all functional signatures before and on anti-CD20 therapy. B) Correlation of established inflammation and T cell gene signatures with our TIPB signature.

### Supplementary Methods

#### FACS Gating Strategy

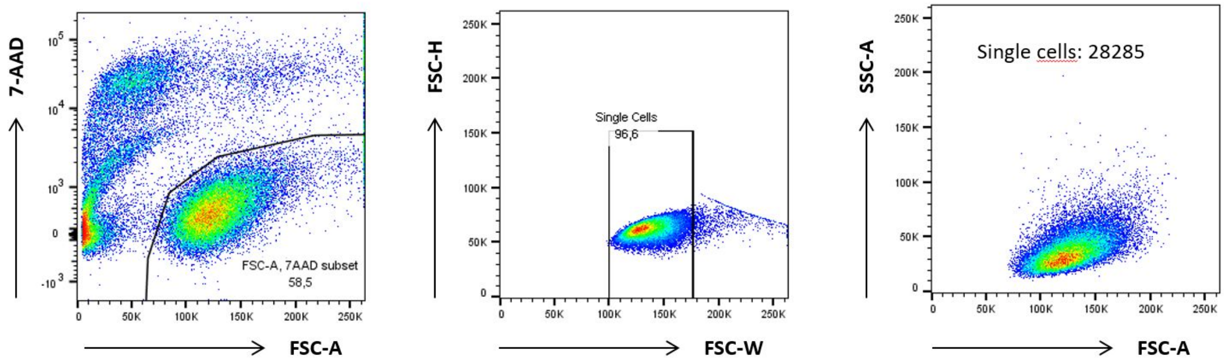


FACS staining of immortalized peripheral blood- and tumor-derived B cells: gating strategy for analysis of viable, single cells.

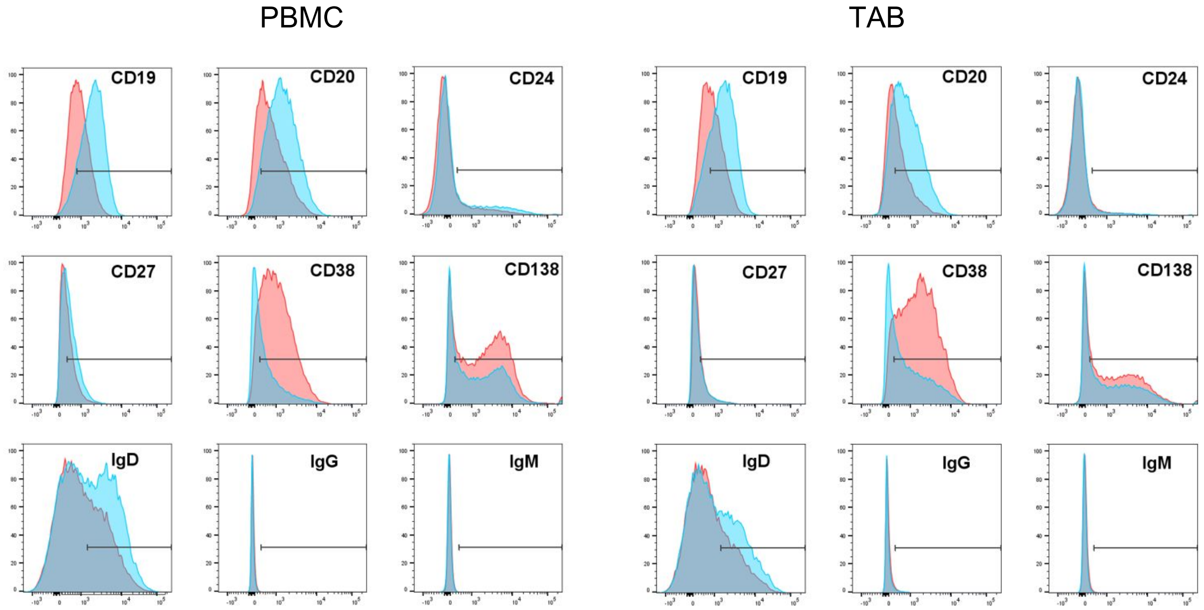


Representative FACS stainings of peripheral blood-derived immortalized B cells (PBMC) and tumor-derived B cells (TAB) of one patient stimulated with control medium (blue histograms) or melanoma-conditioned medium (MCM) (red histograms) for 48 hours. The gate represents the isotype defined positivity of the stainings.
